# Engineering temporal alignment of mRNA delivery enables functional rescue across genetic infertility models

**DOI:** 10.1101/2025.11.13.688269

**Authors:** Qi Jiang, Kexin Su, Guangnan Li, Huali Luo, Hao Wang, Jingwen Luo, Yukai Jason Zhong, Qiaodan Li, Zhihui Zhang, Haifeng Zhang, Wenlin Li, Banghua He, Chenyu Pei, Qin Li, Liang Xu Ma, Hongxiao Cui, Jiaoyan Ma, Deivendran Rengaraj, Linlin Zhang, Xianguang Yang, Zhefan Yuan, Lejun Li, Shuai Liu, Xin Zhiguo Li

## Abstract

Genetic male infertility arises from diverse mutations that disrupt spermatogenesis, yet therapeutic strategies capable of restoring germ-cell function remain limited. Messenger RNA (mRNA) therapeutics offer a programmable and non-integrating approach, but their functional compatibility in structured and immune-privileged tissues remains unclear. Here we identify a temporal mismatch between mRNA delivery kinetics and the progression of spermatogenesis as a key barrier to functional rescue. We show that delivery efficiency does not predict outcome: formulations with high peak expression but short duration fail to support spermiogenesis, whereas sustained expression aligned with developmental timing enables recovery. Guided by this principle, we establish a testis-targeted mRNA delivery platform that restores spermatogenesis across multiple genetic models and generates functional sperm capable of supporting embryo development and multigenerational inheritance. The approach is modular, extends to human seminiferous tubules *ex vivo*, and is supported by comprehensive safety analyses. Together, these findings define a design constraint for RNA therapeutics in structured tissues: functional rescue requires temporal alignment between delivery kinetics and the therapeutic window defined by tissue-specific biological programs.

---

Genetic male infertility affects a substantial fraction of reproductive-age men worldwide and arises from highly heterogeneous defects in spermatogenesis^1,2^. Although assisted reproductive technologies such as intracytoplasmic sperm injection (ICSI) can bypass certain functional deficits, they remain ineffective in the absence of viable sperm and do not address the underlying molecular causes of disease^3,4^. These limitations highlight the need for therapeutic strategies capable of restoring endogenous spermatogenesis.

Messenger RNA (mRNA) therapeutics provide a programmable and non-integrating approach for transient protein replacement and have shown remarkable success in hepatic and intramuscular delivery contexts. However, extending mRNA therapeutics to structured and immune-privileged tissues remains a major challenge. The testis represents a particularly stringent example: the blood–testis barrier (BTB) restricts molecular access, and germ-cell development is tightly coordinated across spatially and temporally organized stages^5–7^. In such systems, it remains unclear which delivery parameters determine whether restored expression translates into functional biological outcomes.

Recent studies have demonstrated proof-of-concept mRNA delivery to the testis and partial restoration of spermatogenesis in specific genetic models^8,9^. However, these approaches have largely focused on delivery efficiency or peak expression as primary metrics of success. Notably, restoration of spermatogenic markers does not consistently result in functional fertility^10,11^, suggesting that additional, unresolved constraints govern the relationship between mRNA delivery and biological function. More broadly, why delivery efficiency alone is insufficient to predict therapeutic outcome in temporally structured tissues remains an open question.

Here, we identify a temporal constraint underlying mRNA-based rescue of spermatogenesis. We show that delivery efficiency does not predict functional outcome, and instead demonstrate that successful rescue requires temporal alignment between mRNA expression kinetics and the biological progression of spermatogenesis. Guided by this principle, we establish a modular mRNA–lipid nanoparticle (LNP) platform that restores spermatogenesis across multiple genetic infertility models, generates functional sperm capable of supporting embryo development and multigenerational inheritance, and enables efficient mRNA delivery in human seminiferous tissue *ex vivo*. Together, our findings define a design constraint for RNA therapeutics in structured tissues and provide a framework for extending mRNA-based therapies beyond the liver.

## Results

### Efficient intratubular delivery enables broad access to the seminiferous epithelium

Delivering nucleic acids into the immune-privileged testis is challenging owing to the BTB and the highly organized architecture of seminiferous tubules. We first compared non-viral delivery approaches, including electroporation, protamine-mediated transfection and red blood cell membrane (RBCM) vesicle encapsulation. None of these strategies produced appreciable mRNA expression in adult mouse testes, indicating limited tissue penetration and inefficient cytosolic release (Extended Data Fig. 1a,b). In contrast, LNPs enabled robust mRNA delivery. To determine how anatomical access influences delivery efficiency, we compared interstitial injection with intratubular infusion via the efferent duct. Intratubular delivery, which fills seminiferous tubules through the rete testis, yielded 2.3–6.1-fold higher luciferase activity than interstitial administration across 6–24 hours (*P* < 0.01; Fig. 1a). This approach was well tolerated and required lower doses than systemic delivery, and was therefore used for subsequent experiments.

**Figure 1.**
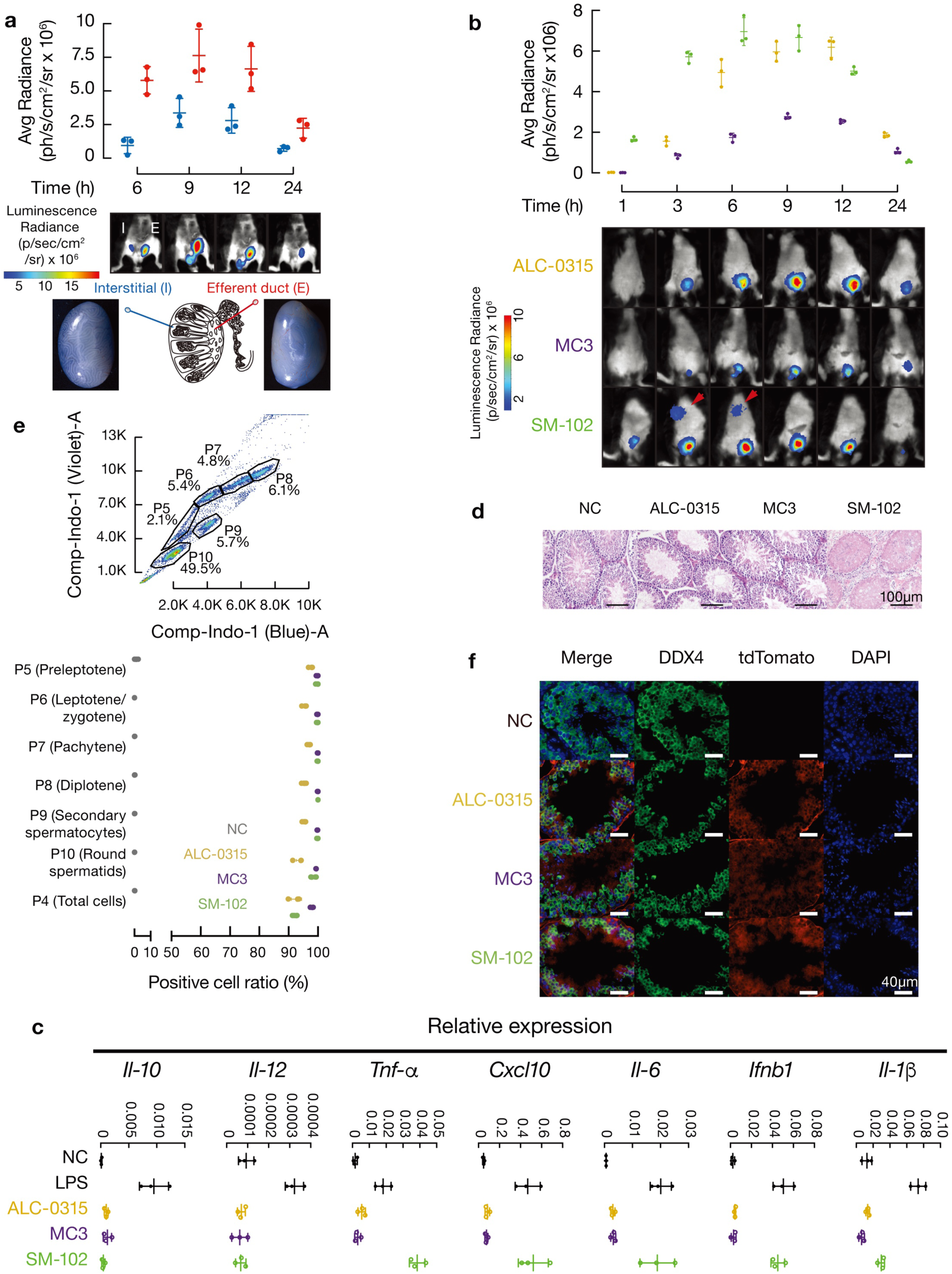
Efficient intratubular mRNA delivery enables seminiferous epithelium access and reveals context-dependent biodistribution and inflammation. (a) Bioluminescence imaging comparing luciferase expression after intratubular (efferent duct, E) versus interstitial (I) injection within the same mouse. Quantification of photon flux at indicated time points (mean ± s.d.; n = 3 mice). Two-way ANOVA with Sidak’s correction. (b) Time course of testicular luciferase expression following intratubular delivery of MC3-, ALC-0315- or SM-102-formulated mRNA. Representative images (1 h and 24 h) and quantification (mean ± s.d.; n = 3 mice). Two-way ANOVA with Sidak’s correction. (c) Innate immune response in testis 12 h after intratubular delivery, measured by RT–qPCR for *Il-10*, *Il-12b*, *Tnf-α*, *Cxcl10*, *Il-6*, *Ifnb1* and *Il-1*β. LPS, lipopolysaccharide. Mean ± s.d. (n = 3 mice). One-way ANOVA with Tukey’s correction. (d) Representative H&E staining 12 h after intratubular injection. SM-102 induces interstitial edema and leukocyte infiltration; MC3 and ALC-0315 show minimal inflammation. Scale bar, 100 µm. (e) Flow cytometric analysis of germ-cell populations (1N, 2N, 4N) and percentage of tdTomato-positive cells following Cre mRNA delivery. Mean ± s.d. (n = 6 testes). One-way ANOVA with Tukey’s correction. (f) Immunofluorescence of Ai9 reporter testes showing tdTomato (red) and DDX4 (green). Nuclei counterstained with DAPI (blue). Scale bar, 40 µm.

We next evaluated three clinically relevant ionizable lipids—MC3, ALC-0315 and SM-102—formulated under comparable physicochemical conditions (Extended Data Fig. 1c,d). All formulations exhibited similar particle sizes (∼100 nm), low polydispersity and high encapsulation efficiency, enabling direct comparison of their *in vivo* performance. Following intratubular administration, SM-102 and ALC-0315 produced higher peak luciferase signals than MC3 (Fig. 1b). However, SM-102 showed prominent off-target accumulation in liver and spleen, even at reduced doses, and induced marked inflammatory responses in the testis (Fig. 1c,d, Extended Data Fig. 1e-l). In contrast, MC3 and ALC-0315 exhibited minimal systemic leakage and limited immune activation. Flow cytometry and immunofluorescence analyses further demonstrated that both MC3 and ALC-0315 achieved efficient transfection across germ-cell populations and Sertoli cells, indicating broad access to the seminiferous epithelium (Fig. 1e,f; Extended Data Fig. 1m,n,o), rather than exhibiting strict cell-type specificity.

These findings establish intratubular LNP delivery as an efficient strategy for accessing germ-cell–containing compartments and reveal formulation-dependent differences in tissue retention, biodistribution and immunogenicity within the seminiferous environment. We further evaluated the impact of RNA format and modification on expression kinetics and immune activation. Among the formats tested, N¹-methylpseudouridine (m^1^Ψ)–modified linear mRNA provided the most favorable balance of robust expression and minimal immune response, whereas self-amplifying and circular RNA exhibited lower expression and increased innate immune activation (Extended Data Fig. 2a-g). We therefore selected modified linear mRNA for subsequent experiments.

### Sustained mRNA expression in the seminiferous epithelium reveals a temporal constraint

To investigate whether differences in delivery behavior influence biological outcomes, we next examined the expression kinetics of mRNA delivered by distinct LNP formulations. We selected *Papolb* (also known as Tpap) as a model target, as it encodes a testis-specific cytoplasmic poly(A) polymerase required for spermiogenesis. Loss of *Papolb* results in a well-characterized arrest at the round spermatid stage without affecting earlier spermatogenic processes^12^, providing a tractable system to assess whether transient protein restoration can drive post-meiotic differentiation.

We engineered an optimized *Papolb* mRNA construct containing the endogenous 5′UTR, a codon-optimized coding sequence, and a stabilizing human α-globin 3′UTR (Fig. 2a). RNA quality control confirmed high integrity, minimal double-stranded RNA contamination, and low endotoxin levels (Extended Data Fig. 3a,b). Following intratubular delivery in wild-type mice, both MC3- and ALC-0315-formulated LNPs mediated robust HA–PAPOLB protein expression within 10 hours (Fig. 2b,c). Expression was restricted to the injected testis and not detected in other organs (Extended Data Fig. 3c), and tissue morphology remained normal (Extended Data Fig. 3d), indicating that exogenous PAPOLB expression is well tolerated. Despite similar initial expression, the persistence of protein and mRNA differed markedly between formulations. MC3-mediated protein expression remained detectable for approximately 10 days by immunofluorescence, whereas ALC-0315-mediated expression declined to near background within ∼3 days (Fig. 2b). Western blot analysis showed a similar trend, with protein detectable for ∼4 days with MC3 and ∼2 days with ALC-0315 (Fig. 2c). At the transcript level, MC3-delivered *Papolb* mRNA persisted for up to ∼14 days, whereas ALC-0315-delivered transcripts became undetectable by ∼5 days (Fig. 2d).

**Figure 2.**
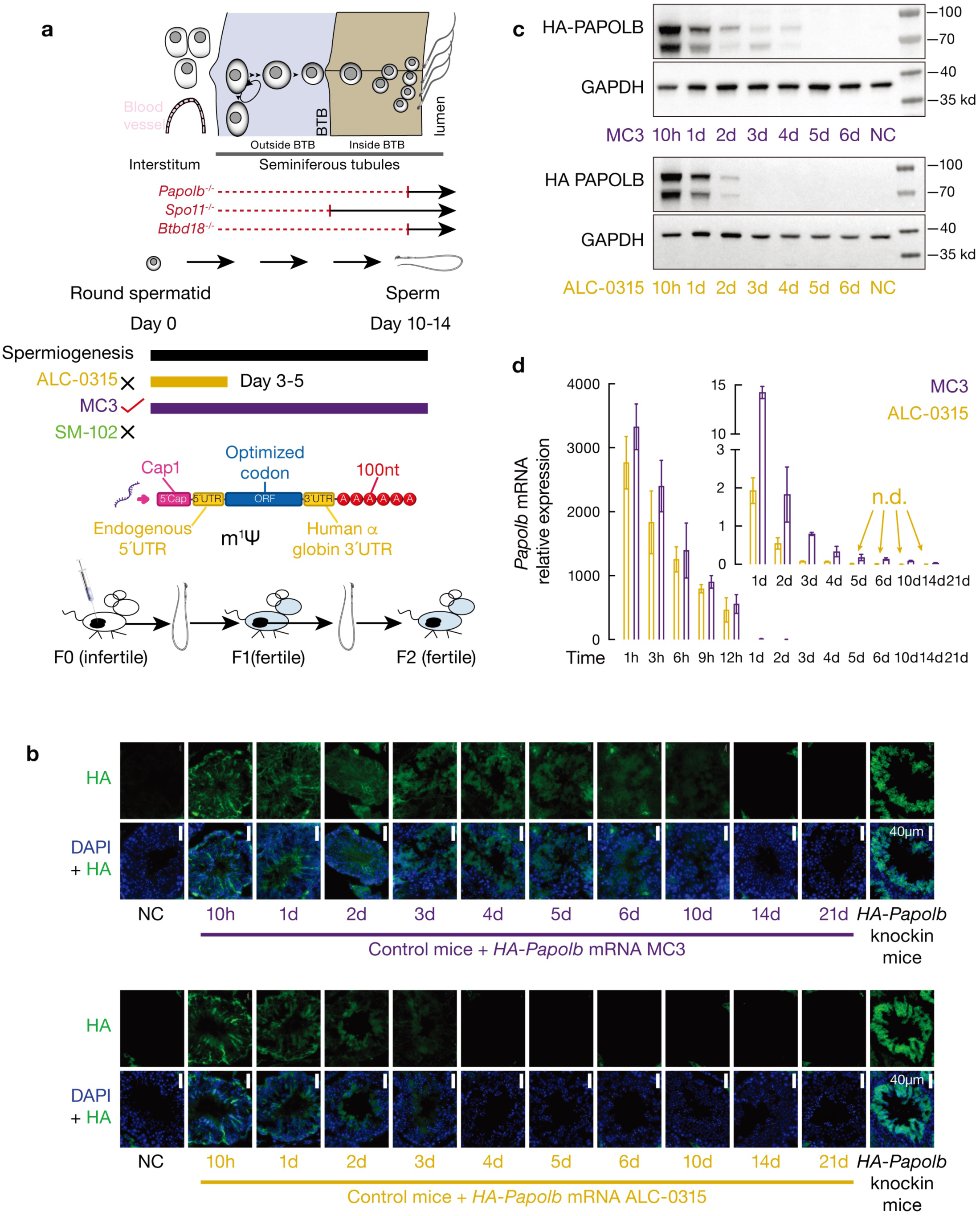
Differential expression kinetics of Papolb mRNA delivered by MC3 and ALC-0315 LNPs. (a) Conceptual framework for mRNA delivery in the seminiferous epithelium and its temporal alignment with spermiogenesis. Schematic illustrating the anatomical organization of the testis, including the blood–testis barrier (BTB), and the developmental progression of germ cells from round spermatids to mature spermatozoa. Genetic infertility models (*Papolb^⁻/⁻^*, *Spo11^⁻/⁻^* and *Btbd18^⁻/⁻^)* arrest at distinct stages of spermatogenesis. The mRNA construct used in this study contains an endogenous 5’UTR, codon-optimized coding sequence and human α-globin 3’UTR with N¹-methylpseudouridine (m¹Ψ) modification. Intratubular delivery of mRNA–LNPs enables transient protein expression to restore spermatogenesis and generate fertile offspring across generations. Expression duration differs across LNP formulations relative to the spermiogenic window (∼10–14 days): ALC-0315 mediates short-lived expression (∼3–5 days) that does not span the required developmental interval, whereas MC3 provides sustained expression that overlaps with the spermiogenic window, enabling complete differentiation. (b) Immunofluorescence detection of HA–PAPOLB expression 10 h after intratubular delivery. Signal is restricted to injected testes. Scale bar, 40 µm. (c) Western blot time course of HA–PAPOLB expression following delivery by MC3 or ALC-0315. GAPDH serves as loading control. (d) RT–qPCR quantification of Papolb mRNA persistence. MC3 exhibits prolonged transcript retention compared with ALC-0315. Mean ± s.d. (n = 3 mice). n.d.: not detected. Two-way ANOVA with Sidak’s correction.

Thus, LNP formulations exhibit a trade-off between peak expression and expression duration: ALC-0315 achieves higher initial expression but short persistence, whereas MC3 produces more sustained expression with lower peak intensity. These differences in delivery kinetics and functional outcomes are summarized in Extended Data Fig. 1p. Given that spermiogenesis from round spermatids to elongating spermatids requires approximately 10–14 days in the mouse testis^13^, this trade-off introduces a temporal constraint, raising the question of which expression profile is compatible with functional differentiation.

### Temporally aligned mRNA delivery enables functional rescue of spermiogenesis

To determine which expression profile supports functional differentiation, we delivered *Papolb* mRNA to *Papolb* knockout mice. A single intratubular injection of MC3-formulated *Papolb* mRNA restored spermatid differentiation, as evidenced by the appearance of elongating spermatids by day 7 (Fig. 3a). In contrast, ALC-0315–formulated *Papolb* mRNA failed to restore spermiogenesis at any examined time point despite achieving higher early expression (Fig. 3a; Extended Data Fig. 4a).

**Figure 3.**
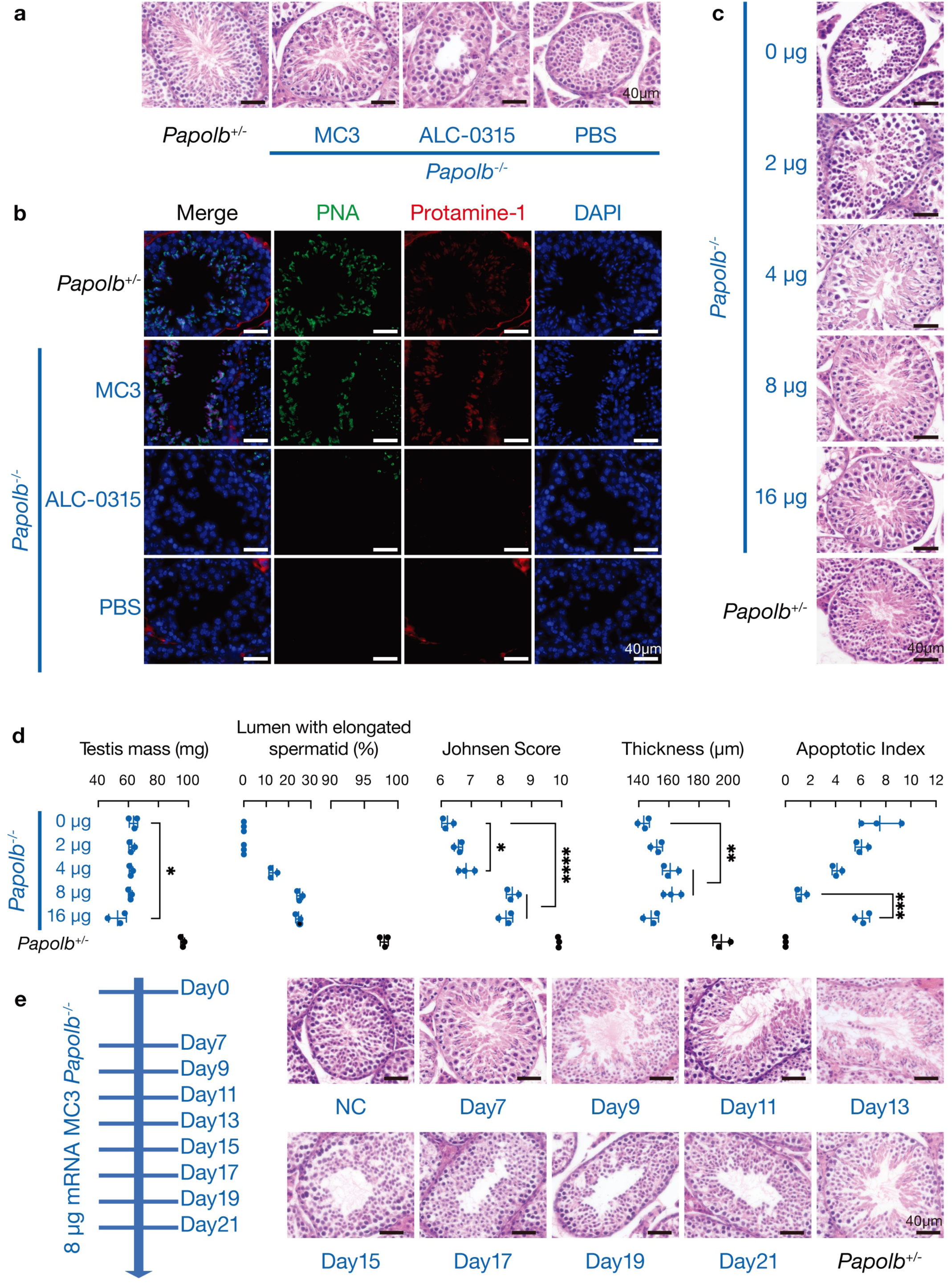
Temporally aligned mRNA delivery restores spermiogenesis in *Papolb*-deficient testes. (a) H&E staining showing restoration of elongating spermatids in MC3-treated *Papolb^−/−^* testes. Scale bar, 40 µm. (b) Immunofluorescence staining for Protamine-1 (red) and acrosomes (PNA, green). Nuclei counterstained with DAPI (blue). Scale bar, 40 µm. (c) Dose-dependent rescue of spermatogenesis across *Papolb* mRNA concentrations. Scale bar, 40 µm. (d) Quantification of spermatogenic recovery, including testis mass, percentage of tubules with elongating spermatids, Johnsen score, epithelial thickness and apoptosis index. Mean ± s.d. (n = 3 mice). One-way ANOVA with Tukey’s correction. (e) Time course of spermatogenic recovery following a single MC3–*Papolb* mRNA injection. Scale bar, 40 µm.

Restoration of post-meiotic progression in MC3-treated testes was confirmed using stage-specific markers. Protamine-1 and transition protein 1 (TNP1) staining demonstrated appropriate nuclear condensation during elongation, ACRV1 and PNA staining confirmed acrosome formation (Fig. 3b; Extended Data Fig. 4b,c). In contrast, markers of earlier spermatogenic stages, including SYCP1, SYCP3, and γH2AX, remained unchanged, indicating that rescue selectively restores post-meiotic differentiation without perturbing upstream processes (Extended Data Fig. 4c,d). Spermatogonial proliferation (PCNA) and global chromatin organization (H3) were also preserved (Extended Data Fig. 4e).

Dose–response analysis revealed that approximately 8 μg *Papolb* mRNA per testis maximally rescued ∼25% of seminiferous tubules (Fig. 3c,d). Lower doses resulted in weaker rescue, whereas higher doses increased apoptosis without further improving recovery (Fig. 3c,d; Extended Data Fig. 4f-h), indicating a defined therapeutic window in which sufficient expression supports differentiation while excessive dosing introduces tissue stress.

Temporal analysis further revealed that elongating spermatids appeared by day 7, peaked between days 9–13, and declined thereafter, with tubules reverting to the arrest phenotype by day 17 (Fig. 3e; Extended Data Fig. 4i-k). This transient wave closely matches the seminiferous epithelial cycle (∼207 h)^14^, indicating that rescued germ cells progress through a single developmental cycle following transient mRNA delivery. Repeated dosing extended the duration of sperm production but did not increase the proportion of rescued tubules (Extended Data Fig. 4l), suggesting that rescue is constrained by the developmental state of individual tubules at the time of treatment.

Together, these findings demonstrate that sustained expression, rather than peak delivery efficiency, is required to support spermiogenesis, consistent with the need to maintain protein expression across the developmental interval of post-meiotic differentiation.

### Rescued spermatogenesis produces functionally competent sperm

Having established that sustained mRNA expression enables morphological rescue of spermiogenesis, we next asked whether the resulting germ cells are functionally competent. Because intratubular delivery disrupts epididymal sperm transport (Extended Data Fig. 5a–c), we assessed sperm function using testicular sperm. Sperm recovered from MC3-treated *Papolb*-deficient testes exhibited normal head–tail morphology comparable to heterozygous controls (Fig. 4a,b). Mitochondrial membrane potential, assessed using JC-1, TMRE and Mitotracker assays, showed preserved polarization patterns similar to controls (Fig. 4c; Extended Data Fig. 5d,e), indicating intact bioenergetic function. DNA integrity, evaluated by alkaline comet assay, revealed no increase in fragmentation relative to control sperm (Fig. 4d). Together, these analyses demonstrate that rescued sperm are structurally intact and metabolically competent.

**Figure 4.**
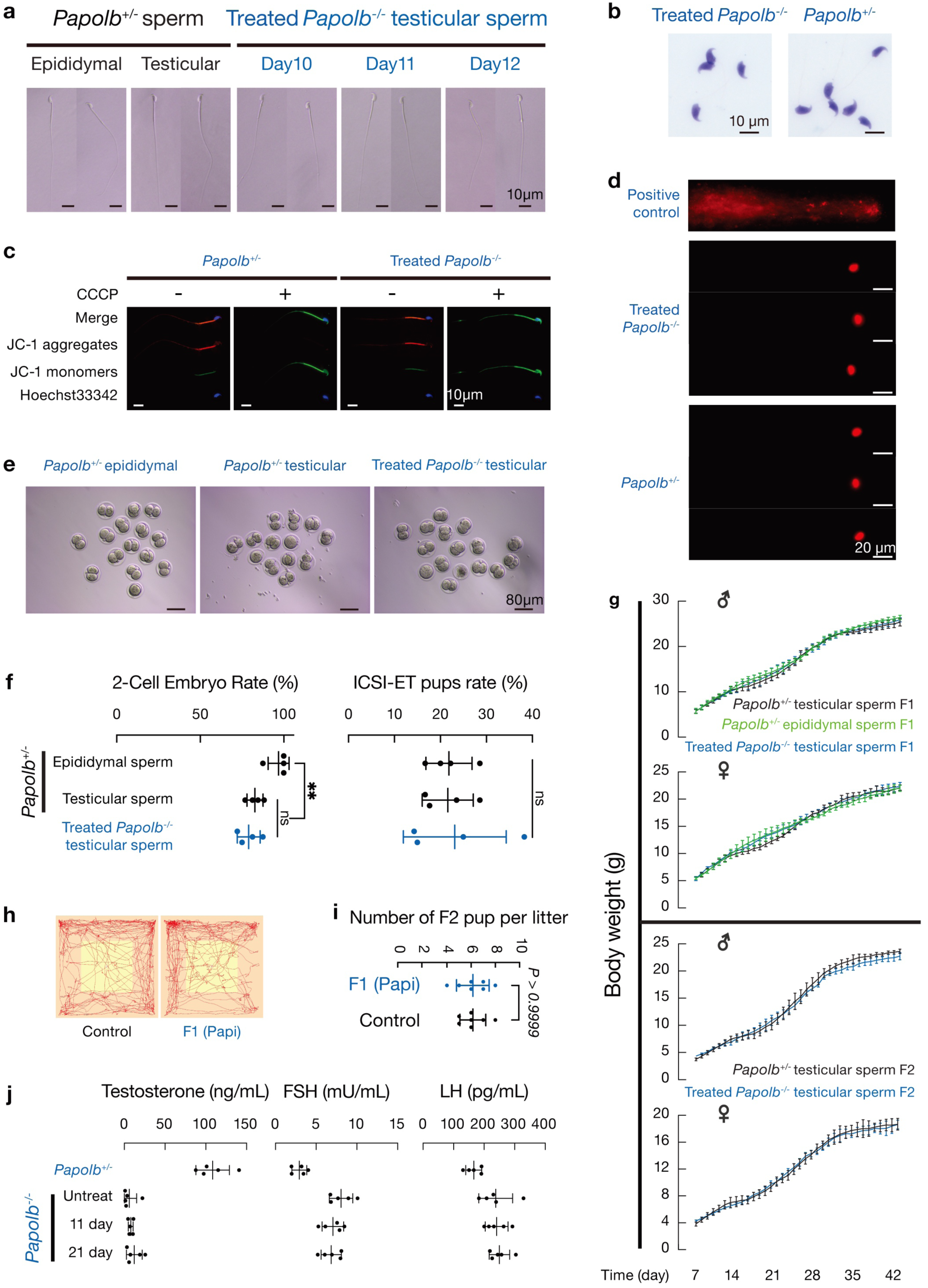
Rescued spermatogenesis produces functionally competent sperm and supports germline transmission. **(a)** Bright-field images of sperm isolated from testes ∼10–12 days post-treatment. Left: epididymal sperm; middle: testicular sperm from *Papolb^+/−^* control; right: testicular sperm from MC3-treated *Papolb^−/−^* male. Scale bar, 10 µm. (n = 3 mice). **(b)** Sperm morphology assessed by Diff-Quik staining comparing testicular sperm from treated *Papolb^−/−^* mice and *Papolb^+/−^* controls. Scale bar, 10 µm. (n = 3 mice per group; ≥100 sperm scored per mouse). **(c)** Mitochondrial membrane potential assessed by JC-1 staining; CCCP-treated sperm served as depolarization control. Scale bar, 10 µm. (mean ± s.d.; n = 3 mice per group; 200 sperm per mouse). CCCP, carbonyl cyanide m-chlorophenyl hydrazone. **(d)** DNA integrity assessed by alkaline comet assay; H₂O₂-treated sperm served as positive control. Scale bar, 20 µm. (mean ± s.d.; n = 3 mice per group; 100 sperm per mouse). **(e)** Representative two-cell embryos 24 h after ICSI using different sperm sources. Scale bar, 80 µm. **(f)** Left, two-cell embryo formation rate (percentage of injected metaphase II oocytes developing to two-cell embryos). Right, live birth rate (percentage of transferred embryos resulting in pups). Data represent mean ± s.d. from four independent experiments (n = 4 ICSI sessions). Statistical analysis: one-way ANOVA with Tukey’s multiple comparisons test. **(g)** Growth curves (days 7–42) for male and female offspring generated via ICSI (F1) and via natural mating (F2). Data are mean ± s.d. (n = 5 per sex per group). Statistical analysis: two-way ANOVA with Tukey’s multiple comparisons test. **(h)** Representative locomotor tracks from open-field testing (n = 5 mice per group). **(i)** Pups per litter for F1 offspring derived from treated *Papolb^−/−^* sires versus *Papolb^+/−^* controls. Data are mean ± s.d. (n = 7 litters per group). Statistical analysis: unpaired t-test (two-sided). **(j)** Serum reproductive hormone levels in treated F0 mice. Testosterone, FSH, and LH were measured by ELISA in *Papolb^+/−^* controls, untreated *Papolb^−/−^*(NC), and treated *Papolb^−/−^* mice at 11 days and 6 months post-treatment. Samples were collected at 9:00–10:00 am. Data are mean ± s.d. (n = 5 mice per group). Statistical analysis: one-way ANOVA with Tukey’s multiple comparisons test. FSH, follicle-stimulating hormone; LH, luteinizing hormone.

We next evaluated developmental competence using intracytoplasmic sperm injection (ICSI). Testicular sperm from MC3-treated *Papolb*-deficient mice supported embryo development at rates comparable to heterozygous controls, with ∼79% of injected oocytes progressing to the two-cell stage (Fig. 4e,f). Transfer of two-cell embryos yielded live F1 offspring, and embryos progressed through normal cleavage stages *in vitro* (Extended Data Fig. 5f). Offspring derived from rescued sperm exhibited normal growth trajectories, morphology and behavior (Fig. 4g-h; Extended Data Fig. 5g–m; Table S1). Sex ratios were comparable across groups (Extended Data Fig. 5n), and no developmental abnormalities were observed. Importantly, F1 males were fertile and produced F2 and F3 generations through natural mating (Fig. 4i; Extended Data Fig. 5o,p), demonstrating stable germline transmission. All offspring carried one mutant and one wild-type allele, consistent with transient mRNA supplementation rather than genomic modification (Extended Data Fig. 5q).

To assess endocrine consequences, we measured serum testosterone, FSH and LH levels. As expected, untreated *Papolb* mutants exhibited reduced testosterone and compensatory elevation of gonadotropins. In contrast, hormone levels in F1 and F2 progeny fell within normal physiological ranges (Fig. 4j; Extended Data Fig. 5r), indicating preserved endocrine homeostasis across generations.

Together, these results demonstrate that temporally aligned mRNA delivery restores spermatogenesis to a level sufficient for ICSI competence and supports normal development and germline transmission.

### Temporally aligned delivery enables functional rescue across distinct infertility etiologies

We next asked whether the temporal constraint identified above generalizes across distinct genetic causes of spermatogenic failure. To this end, we applied the optimized MC3-based mRNA delivery strategy to *Spo11* and *Btbd18* knockout models, which represent mechanistically distinct defects at meiotic and post-meiotic stages, respectively^15–17^.

In *Spo11*-deficient mice, which arrest at early meiotic stages due to failure of homologous recombination, intratubular delivery of *Spo11* mRNA restored meiotic progression, as demonstrated by chromosome spread analysis showing the emergence of pachytene and diplotene cells (Fig. 5a). In both *Spo11* and *Btbd18* models, elongating spermatids appeared at later time points consistent with their respective developmental blocks (Fig. 5b; Extended Data Fig. 6a-c), indicating that rescue follows intrinsic stage-dependent timing. Quantitative analysis revealed restoration of spermatogenesis in approximately 19.6% of seminiferous tubules in *Spo11* mutants and 12.9% in *Btbd18* mutants, whereas no rescue was observed in control-treated animals (Extended Data Fig. 6d). As in the *Papolb* model, mature sperm accumulation in the epididymis was not observed, consistent with disruption of sperm transport caused by the delivery procedure (Extended Data Fig. 6e).

**Figure 5.**
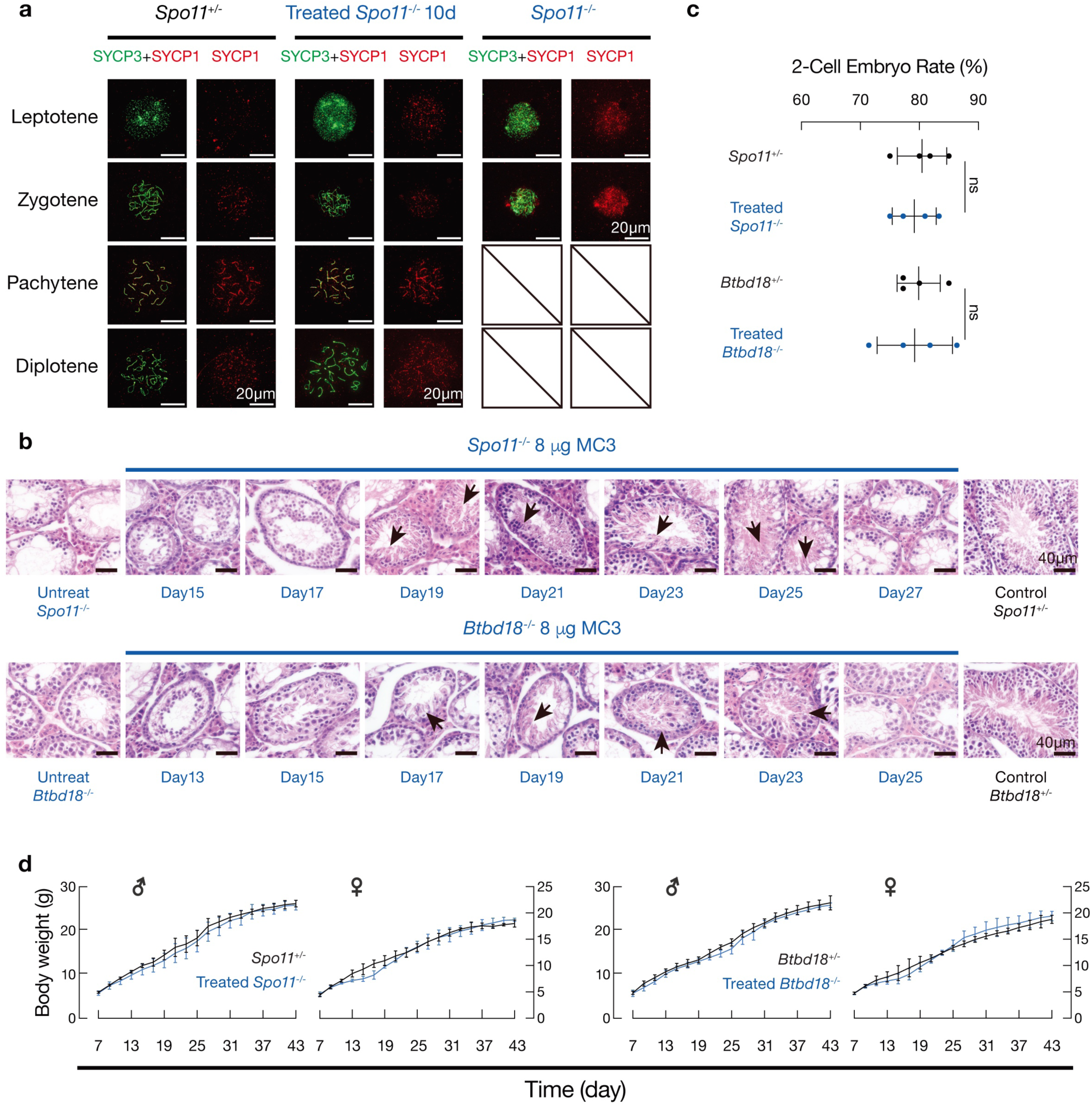
Functional rescue across distinct infertility etiologies. **(a)** Chromosome spread analysis of *Spo11* rescue. SYCP3 (green) and SYCP1 (red) staining. *Spo11^−/−^*cells arrest at early zygotene; treated *Spo11^−/−^* testes show pachytene and diplotene cells. Scale bar, 20 µm. (n = 3 mice per group; 100 nuclei scored per mouse). **(b)** H&E staining showing emergence of elongating spermatids at indicated days post-treatment in *Spo11^−/−^* and *Btbd18^−/−^* testes. Scale bar, 40 µm. (n = 3 mice per group per time point). **(c)** Two-cell embryo formation after ICSI using testicular sperm from treated *Spo11^−/−^* or *Btbd18^−/−^* males compared with heterozygous controls. Scale bar, 80 µm. (n = 4 sessions; ∼20 embryos per group per session). Statistical analysis: unpaired t-test (two-sided). **(d)** Postnatal growth curves for male and female F1 offspring derived by ICSI using sperm from treated *Spo11^−/−^* or *Btbd18^−/−^* males compared with heterozygous controls. Data are mean ± s.d. (n = 5 per sex per group). Statistical analysis: two-way ANOVA with Tukey’s multiple comparisons test.

Functional competence of rescued sperm was confirmed by ICSI. Testicular sperm from treated *Spo11* and *Btbd18* mice supported embryo development and generated viable offspring with normal growth (Fig. 5c,d; Extended Data Fig. 6f–i). Sex ratios remained within expected ranges across models (Extended Data Fig. 6j), indicating no detectable developmental bias. These findings demonstrate that temporally aligned mRNA delivery enables functional rescue across multiple infertility models spanning distinct genetic etiologies and developmental stages, supporting the general applicability of this design principle.

### The mRNA–LNP platform exhibits translational feasibility and long-term safety

Given the germline-adjacent nature of this intervention, we next evaluated safety at multiple levels, including tissue integrity, systemic toxicity, and germline genetic and epigenetic stability. At the tissue level, no evidence of chronic inflammation or fibrosis was observed in treated testes at either early or late time points (Fig. 6a,b). Histological analysis showed preserved seminiferous architecture without tubular atrophy or interstitial remodeling (Extended Data Fig. 7a). Claudin11 staining indicated intact blood–testis barrier (BTB) integrity (Fig. 6c), and CYP17A1 expression demonstrated preserved Leydig cell steroidogenic function (Fig. 6c; Extended Data Fig. 7d). These findings indicate that intratubular mRNA delivery does not disrupt key somatic compartments within the testis.

**Figure 6.**
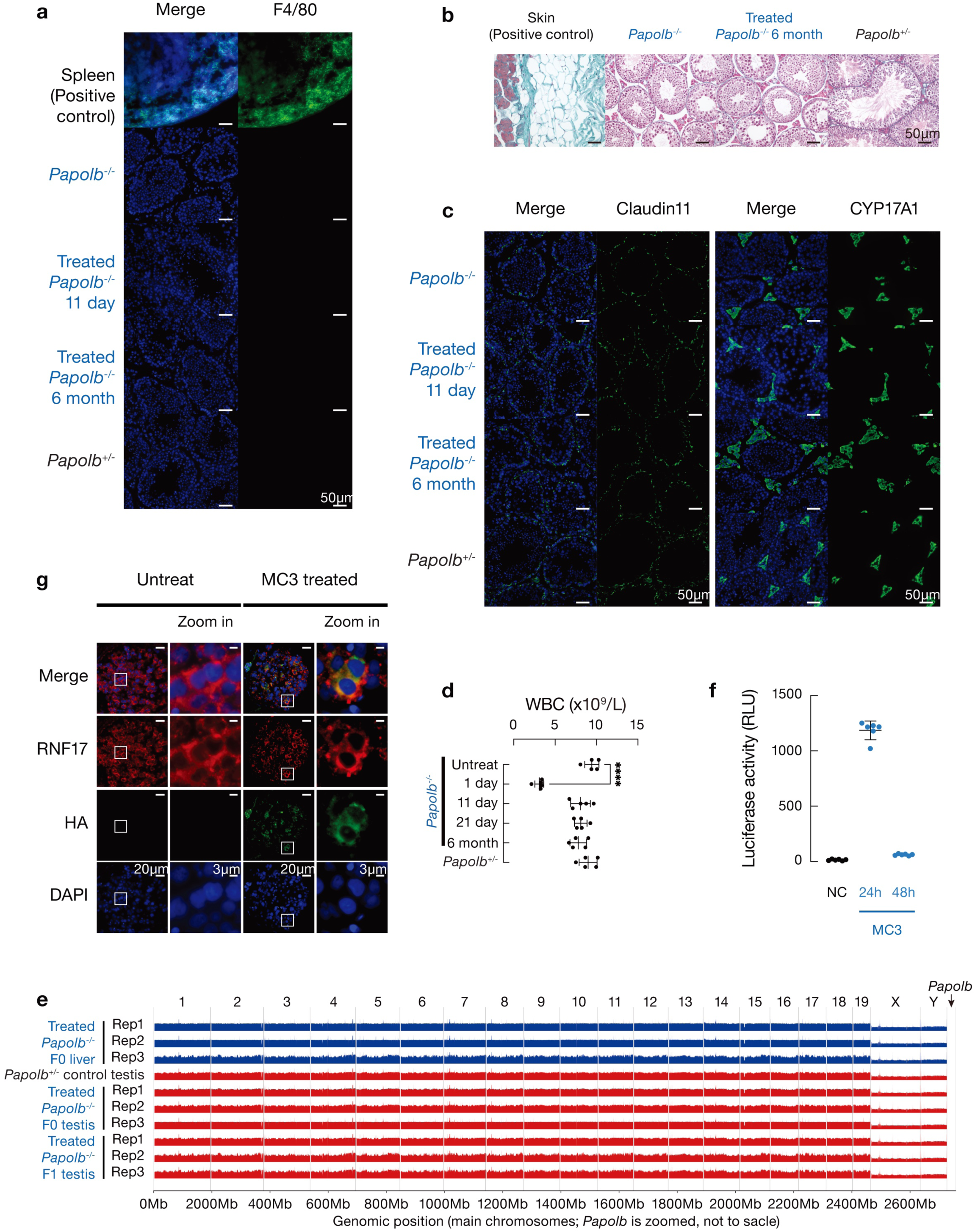
Translational safety and human tissue compatibility of mRNA–LNP delivery. **(a)** F4/80 immunofluorescence analysis of macrophage infiltration in testes at 11 days and 6 months post-treatment. Spleen served as positive control. Scale bar, 50 µm. (n = 3 mice per group; representative of 3 sections per testis). **(b)** Masson’s trichrome staining of testes at 6 months post-treatment. Skin served as positive control for collagen staining. (n = 3 mice per group; representative of 3 sections per testis). **(c)** Claudin11 (left) and CYP17A1 (right) immunostaining of blood–testis barrier integrity in testes at 11 days and 6 months post-treatment. Scale bar, 50 µm. (n = 3 mice per group; representative of 3 sections per testis). **(d)** Systemic safety evaluation by complete blood count (WBC) as representative parameter of systemic safety. Data are mean ± s.d. (n = 5 mice per group per time point). Statistical analysis: one-way ANOVA with Tukey’s multiple comparisons test. WBC, white blood cell count. **(e)** Whole-genome sequencing–based integration analysis. Structural variant detection was performed on reads aligned to a hybrid reference genome containing the mouse genome and the *Papolb* cDNA contig; no junction-supporting breakend events were detected. n = 3 per tissue group; mean coverage ∼30×. CNV analysis is shown in Extended Data Fig. 6i. **(f)** Luciferase activity in cultured human seminiferous tubules at 0, 24, and 48 h after LNP exposure. (n = 2 donors; 3 replicates per donor). Statistical analysis: one-way ANOVA with Tukey’s multiple comparisons test. **(g)** HA and RNF17 co-localization in human tubule cultures. Scale bar, 20 µm (main) and 3 µm (zoom). (n = 3 donors).

Systemic safety assessment revealed no detectable abnormalities in major organs or serum biochemical parameters across time points (Fig. 6d; Extended Data Fig. 7b, c, e). Transient hematological changes observed shortly after injection resolved during follow-up, consistent with procedural stress rather than sustained toxicity.

To assess germline safety, we performed multigenerational epigenetic and genomic analyses. DNA methylation at imprinting control regions (*H19*, *Peg3*, *Snrpn*) was unchanged in offspring derived from treated males (Extended Data Fig. 7f, g), and transcriptomic profiles of early embryos showed high concordance with controls (Extended Data Fig. 7h), indicating preservation of early developmental programs. Transcriptomic analysis of adult tissues from F1 offspring, including heart, kidney, liver, spleen, lung, and testis, similarly showed high concordance with controls, with no consistent or pathway-level alterations detected (Extended Data Fig. 7h), indicating absence of systemic transcriptional perturbation. Whole-genome sequencing at ∼30× coverage revealed no evidence of transgene integration or large-scale structural variation in F0 or F1 samples (Fig. 6e; Extended Data Fig. 7i), and PCR-based assays confirmed the absence of detectable integration events (Extended Data Fig. 7j).

Finally, to evaluate translational feasibility, we examined mRNA delivery in human seminiferous tubule cultures (Extended Data Fig. 8a, b). Efficient expression of reporter mRNA was observed in germ-cell–containing compartments across multiple donors (Fig. 6f; Extended Data Fig. 8c), and delivered protein co-localized with RNF17-positive germ cells (Fig. 6g; Extended Data Fig. 8d). These results demonstrate that the optimized mRNA–LNP platform is compatible with human testicular tissue under *ex vivo* conditions.

Collectively, these findings indicate that temporally aligned mRNA delivery can be achieved without detectable long-term toxicity, genomic perturbation or epigenetic disruption, and supports translational potential in human tissue.

## Discussion

Messenger RNA therapeutics have rapidly advanced as a platform for *in vivo* protein replacement, yet their extension beyond the liver remains a central challenge in the field. In particular, delivery to structured and developmentally dynamic tissues has remained difficult, as conventional metrics such as peak transfection efficiency do not necessarily translate into functional outcomes. Here, we identify a fundamental constraint underlying this limitation: the temporal alignment between mRNA expression kinetics and the biological progression of the target tissue.

Our findings reveal that delivery efficiency alone is insufficient to predict functional rescue in the seminiferous epithelium. Formulations such as ALC-0315 and SM-102 achieve high initial expression but exhibit short persistence or induce tissue perturbation, limiting their functional compatibility. In contrast, MC3-mediated delivery produces sustained expression that overlaps with the developmental window of spermiogenesis, enabling completion of post-meiotic differentiation. These observations demonstrate that therapeutic efficacy is governed not by maximal expression, but by whether expression duration matches the intrinsic timescale of the biological process being targeted.

This principle provides a conceptual framework for interpreting prior observations in the field. Previous studies have reported restoration of spermatogenic morphology following mRNA delivery without recovery of functional fertility^10,11^, suggesting that expression of differentiation markers does not guarantee developmental completion. Our results provide a unifying explanation for these discrepancies: transient expression that fails to span the full developmental interval may initiate differentiation but cannot sustain its completion. By contrast, temporally matched expression enables a full differentiation trajectory sufficient for functional sperm production.

Beyond a single genetic model, we demonstrate that this temporal constraint applies across distinct infertility etiologies. Using *Spo11* and *Btbd18* models, which represent mechanistically different defects at meiotic and post-meiotic stages, we show that temporally aligned mRNA delivery restores spermatogenesis and produces functional offspring. These findings indicate that the requirement for temporal alignment is not specific to a particular gene or pathway, but reflects a broader property of spermatogenic progression. More generally, this suggests that RNA therapeutics targeting dynamic biological systems must be designed to match the temporal structure of the underlying process.

From a translational perspective, our study establishes a modular and clinically grounded platform for mRNA-based treatment of genetic male infertility. Unlike previous approaches that rely on newly engineered or proprietary lipid formulations, our strategy leverages clinically validated ionizable lipids, enabling more direct interpretation of safety and scalability. We further develop a comprehensive safety framework spanning long-term tissue integrity, somatic cell function, multigenerational phenotyping, epigenetic stability and genome-wide integration analysis. Across these assessments, we observe no detectable adverse effects within current limits of detection, supporting the feasibility of transient mRNA-based intervention in germline-adjacent tissues.

The translational potential of this approach is further supported by our demonstration of efficient mRNA delivery in human seminiferous tubules *ex vivo*. While the surgical delivery method used in mice disrupts epididymal sperm transport, the anatomical accessibility of the human testis may permit minimally invasive delivery strategies that preserve physiological sperm maturation. Under such conditions, temporally aligned mRNA delivery could potentially enable broader reproductive options, ranging from assisted reproduction to natural conception, depending on the extent of restored spermatogenesis. Further studies in large-animal models will be required to evaluate delivery strategies, dosing paradigms and long-term safety in clinically relevant settings.

Our findings also have broader implications for the design of extrahepatic RNA therapeutics. Many target tissues—including the testis, central nervous system, and regenerating epithelia—are characterized by tightly regulated temporal programs. In such contexts, maximizing delivery efficiency or peak expression may be insufficient or even counterproductive. Instead, therapeutic success may depend on matching the duration and dynamics of expression to the temporal requirements of the target system. We therefore propose temporal alignment as a general design constraint for RNA therapeutics in structured tissues beyond the liver.

In summary, we establish a modular mRNA–LNP platform that restores spermatogenesis across multiple genetic etiologies while maintaining long-term and multigenerational safety. More broadly, we define a principle linking delivery kinetics to biological function, providing a conceptual framework for designing RNA therapeutics in temporally structured systems.

## Methods

### Nanoparticle formulation and characterization

#### Formulation

Lipid nanoparticle (LNP) formulations were prepared by a rapid ethanol dilution method, similar to published protocols^18^. Briefly, an ionizable lipid (such as ALC-0315, SM-102, or MC3; obtained from UTgene, Cat# UT1001, UT1002, UT1003, respectively) and helper lipids – cholesterol (Sigma–Aldrich, C8667), 1,2-distearoyl-sn-glycero-3-phosphocholine (DSPC; Avanti, 850365P), ALC-0159 (AVT (Shanghai) Pharmaceutical Tech, Cat# 002009), and a polyethylene glycol conjugated lipid (PEG-lipid; AVT, Cat# 002005) – were dissolved together in absolute ethanol at defined molar ratios. MC3 LNPs were formulated with DLin-MC3-DMA, DSPC, cholesterol and DMG-PEG2000 at a molar ratio of 50:10:38.5:1.5. For SM-102 LNPs, the molar ratio of SM-102/DSPC/cholesterol/DMG-PEG2000 was 50:10:38.5:1.5. ALC-0315 LNPs were formulated with ALC-0315/DSPC/cholesterol/ALC-0159 at a molar ratio of 46.3:9.4:42.7:1.6. The nucleic acid cargo (e.g., mRNA) was diluted in an acidic buffer (10 mM citrate, pH ∼4.0). The ethanol lipid solution was rapidly mixed with the aqueous RNA solution at a fixed 1:3 volume ratio (organic: aqueous) to promote spontaneous self-assembly of nanoparticles. The mixture was incubated at room temperature for ∼10 minutes to allow vesicle formation. For *in vivo* use, the crude LNP suspension was further purified by dialysis against PBS to remove residual ethanol and any unencapsulated RNA, then concentrated in PBS by ultrafiltration to the desired nucleic acid concentration. The specific ionizable lipid and type of nucleic acid cargo used in each experiment are detailed in the Results (and figure legends). In general, the nucleic acid cargos included various mRNA constructs, a self-amplifying mRNA (saRNA), or a circular RNA (circRNA). The sources of these constructs were as follows: *Papolb* mRNA and saRNA were synthesized in-house by *in vitro* transcription; Cre mRNA was purchased from RNAlfa Biotech (Cat# Alfa-2501); firefly luciferase mRNA was from Levostar (Cat# LS-RM021); and the circRNA was obtained from Novoprotein (Cat# MR201-U100).

#### Characterization

The hydrodynamic diameter and polydispersity index (PDI) of LNPs were measured by dynamic light scattering (DLS) using a Malvern Zetasizer Nano instrument. Samples were diluted in PBS and equilibrated at room temperature before analysis. Size is reported as the intensity-weighted mean diameter, with PDI as a measure of dispersity.

The surface ζ-potential of the nanoparticles was determined on the same instrument using folded capillary cells, after diluting samples ∼50-fold in deionized water.

Encapsulation efficiency of mRNA was assessed with a RiboGreen RNA fluorescence assay (Thermo Fisher Quant-iT™ RiboGreen kit), by treating aliquots of the LNP with or without 0.2% Triton X-100 to distinguish encapsulated vs. total RNA. Encapsulation efficiency (%) was calculated as 100 × (protected RNA / total RNA input).

### Animal experiments

All animal procedures were performed in accordance with institutional and national guidelines for animal welfare and were approved by the Laboratory Animal Welfare and Ethics Review Committee of Zhejiang University (ZJU20220156 and ZJU20250897). The protocol adhered to principles of humane care, and measures were taken to ensure animal well-being throughout the experiments. Housing conditions: temperature 20–26°C, humidity 40–70%, 12 h light/dark cycle (lights on at 7:30; off at 19:30), normal chow diet, corn cob bedding, cage density of 5 mice per cage.

Mouse strains: Wild-type C57BL/6J mice (Jackson Laboratory, stock #000664) were used for most *in vivo* experiments. B6.Cg-*Gt(ROSA)26Sor^tm^*^9^*^(CAG-tdTomato)Hze^*/J reporter mice (the Ai9 line; JAX stock #007909), *Papolb* knockout mice (*Papolb^−/−^*) were obtained from RIKEN BioResource Center^12,19^, *Spo11* knockout mice were constructed according to the previous publication^15^, and *Btbd18* knockout strain^17^ were maintained on a C57BL/6 background. Mice were genotyped by PCR on tail DNA (using primers provided by the source repository). To test the potential integration of *Papolb* mRNA, we used primers targeting the coding sequence of the delivered mRNA (which was codon-optimized, distinct from the genomic copy; primer sequences used for all genotyping were provided in Extended Data Table 1). Unless otherwise noted, experimental cohorts consisted of young adult male mice (aged over 6 weeks, average 25 g body weight) to ensure sexual maturity. Mice were randomly assigned to treatment groups, and investigators were blinded to group allocations during outcome assessments wherever possible (see Statistical analysis below). Number of animals per group: n = 3 for LNP delivery, rescue, and fertility; n = 5 for behavioral assays and sexual activity.

### *In vivo* administration of nanoparticles / mRNA

Two delivery routes were used, depending on the target cell population: (1) injection via the rete testis/efferent duct to deliver material into the seminiferous tubules (targeting primarily Sertoli cells and germ cells), and (2) direct interstitial injection into the testis parenchyma (targeting Leydig cells and other interstitial cells). In each case, adult male mice were anesthetized with 2.5% tribromoethanol in PBS (Avertin, i.p., MeilunBio, MA0478) and placed supine on a sterile surgical platform. For the rete testis (tubular) delivery, a small lower abdominal incision (∼1 cm) was made to exteriorize the testis within the tunica. Using a dissecting microscope, the efferent ducts emerging from the testis (near the junction with the epididymis) were identified. A fine pulled-glass micropipette (tip diameter ∼50–80 µm) pre-loaded with the RNA solution (with a trace of 0.05% Fast Green dye added for visibility) was carefully inserted into the lumen of the efferent duct. Approximately 20 µL of solution was slowly injected at 3–5 µL/sec; successful delivery into the seminiferous tubules was confirmed by observing the dye spreading from the rete testis throughout the tubules (visible as faint green coloration filling the testis) (Fig.1a). For interstitial injections, a 30-gauge needle (Hamilton syringe) was inserted directly into the testis parenchyma, and ∼20 µL of solution was delivered into the interstitial space between tubules at the same rate. Care was taken to inject slowly and avoid excessive pressure to prevent tissue rupture or back-leakage. After either type of injection, the abdominal incision was closed with absorbable sutures. Analgesia: Ibuprofen (50–60 mg/kg/day; 10 mL of Children’s Motrin in 500 mL of water) continuously in drinking water for 3 days post-surgery. Mice were monitored during recovery from anesthesia and returned to their home cages once fully ambulatory.

### Dosing regimens

Doses of the nucleic acid cargo were selected based on preliminary studies and prior literature. Ai9 reporter mice received LNP-encapsulated Cre mRNA via tail vein injection at a dose of 10 µg mRNA per mouse (approximately 0.4 mg/kg body weight). For luciferase reporter studies, mice were administered firefly luciferase mRNA–LNP at 10 µg (0.4 mg/kg) (by intramuscular or intratesticular injection) as detailed in the Results section. *Papolb* knockout mice assigned to gene therapy received an i.t. dose of LNPs carrying *Papolb* mRNA at 0.32 mg/kg. In some experiments, self-amplifying mRNA (saRNA) or circular RNA (circRNA) formulations were administered via the same routes; the dosing for saRNA and circRNA (10 µg per testis, 0.4 mg/kg) was comparable to the mRNA doses. Control animals received equivalent volumes of vehicle (PBS). All treatments were given as a single dose unless otherwise noted. No animals were excluded from analysis a priori except in the case of a technical failure (e.g., if an injection was visibly mis-delivered); in this study, no such exclusions were necessary. Body weights at dosing: 25 g. Dose-response experiments: *Papolb* KO mice received 2, 4, 8, or 16 µg *Papolb* mRNA–LNPs (n = 2 mice per dose). Samples were collected on day 7 post-injection. Time-course experiments: *Papolb* KO mice treated with 8 µg *Papolb* mRNA–MC3 LNPs were sampled at 7, 9, 11, 13, 15, 17, 19, and 21 days post-injection. *Btbd18* KO mice were sampled on day 13 and 19, and *Spo11* KO mice on day 15 and 21.

### *In vivo* fluorescence and bioluminescence imaging

#### Fluorescence imaging (tdTomato reporter)

To evaluate nanoparticle biodistribution and transgene expression *in vivo*, mice were imaged using an IVIS Spectrum optical imaging system (BIOSPACE LAB Photon IMAGER Optima) at various time points after treatment. For the *Cre* mRNA delivery studies using Ai9 reporter mice, animals were euthanized, dissected and major organs (including testes, liver, spleen, kidneys, lungs, and heart) harvested for *ex vivo* fluorescence imaging at 48 hours post-injection. tdTomato fluorescence was captured using the appropriate excitation/emission filter settings (Ex 570 nm / Em 620 nm). Identical imaging parameters (illumination intensity, exposure time, binning, etc.) were used for all animals to allow quantitative comparison. Fluorescence intensity was quantified as radiant efficiency within regions of interest (ROIs) using M3Vision® software (BIOSPACE LAB).

#### Bioluminescence imaging (luciferase reporter)

For luciferase experiments, bioluminescence imaging was performed at designated timepoints. To compare different intratesticular injection routes (rete testis vs interstitial), images were collected at 6, 9, 12, and 24 h. To compare different LNP formulations (MC3, ALC-0315, SM-102) and assess SM-102 leakage, images were collected at 1, 3, 6, 9, 12, and 24 h post-injection. Mice were injected intraperitoneally with D-luciferin (150 mg/kg, 100 µL; Gold Biotechnology, LUCNA-100) and anesthetized with isoflurane 5 min later. Photons emitted from firefly luciferase activity were collected with a 5 s exposure, standard binning, and consistent field of view. Data were analyzed as total photon flux (photons/sec) within ROIs encompassing the testes or other relevant regions. Baseline auto-luminescence was negligible.

Tissue preparation and immunofluorescence microscopy

#### Frozen-section imaging of native tdTomato

For high-resolution analysis, testes and other relevant organs were collected, embedded fresh in optimal cutting temperature (O.C.T.) compound, flash-frozen on dry ice, and stored at −80 °C until sectioning. Cryostat sections (5 µm) were mounted on glass slides, briefly fixed in 4% paraformaldehyde (PFA) or acetone, counterstained with DAPI, and mounted in antifade medium.

#### Immunofluorescence staining of testis sections

Other testis samples were fixed in 4% PFA (overnight at 4 °C), dehydrated, cryoprotected in 30% sucrose, and sectioned at 5 µm. After washing in PBS, sections underwent heat-mediated antigen retrieval (10 min in citrate buffer pH 6.0). Blocking was performed in 5% normal goat serum with 0.1% Triton X-100 for 1 h. Primary antibodies were incubated overnight at 4 °C: SOX9 (HuaBio, Cat# ET1611-56, 1:500), PRM1 (Briar Patch, Hup-1N, 1:200), and HA-tag (CST, clone C29F4, 1:500). Alexa Fluor 488–conjugated peanut agglutinin (PNA, Thermo Fisher L21409; 2 µg/mL) was used for acrosomes. After PBS washes, slides were incubated for 1 h with Alexa Fluor 488 goat anti-rabbit (Invitrogen A-11008, 1:500) and Alexa Fluor 594 goat anti-mouse (Invitrogen A-11005, 1:500). Nuclei were counterstained with DAPI. Imaging was performed on a Zeiss Axio Imager.M2 microscope with fixed acquisition settings across groups.

### Histological analysis

For general assessment of morphology and toxicity, testes were fixed in Bouin’s solution overnight at 4 °C, transferred to 70% ethanol, dehydrated, and paraffin embedded. Sections (3 µm) were stained with hematoxylin and eosin (H&E) using standard protocols. Slides were examined by a blinded observer. Features evaluated included seminiferous tubule structure, presence or absence of elongating/elongated spermatids, epithelial height, and interstitial integrity. Pathological signs such as germ cell degeneration, vacuolization, multinucleated cells, or disrupted tubule organization were recorded. Images were taken for documentation and compared to age-matched controls.

### Intracytoplasmic sperm injection (ICSI) and offspring / fertility analysis

ICSI was performed using testicular sperm from treated males. Mature metaphase II oocytes were obtained from superovulated C57BL/6J females (6–8 weeks) induced with PMSG (NSHF, Cat#110254564) and hCG (NSHF, Cat#A2212051). Cumulus–oocyte complexes were harvested 14 h post-hCG, cumulus cells removed with hyaluronidase (300 µg/mL; Sigma, H4272), and denuded oocytes maintained in KSOM medium (AibeiBio, M1435) under mineral oil (Sigma, M5310-500ML).

Testes from treated males (10–13 days post-injection) were digested with collagenase IV (0.4 mg/mL; Worthington LS004188) and DNase I (10 µg/mL; Roche 10104159001), followed by trypsin-EDTA (0.25%; Gibco 25200072). Digestion was quenched with FBS. Suspensions were filtered, centrifuged, and resuspended in GBSS + 5% FBS + DNase I. Only morphologically normal motile or twitching spermatozoa were selected.

ICSI was performed essentially as described^20,21^ using a piezo-electric manipulator (Eppendorf PiezoXpert). Individual sperm were drawn tail-first into injection pipettes (6.5–8.5 µm ID) containing polyvinylpyrrolidone (Vitrolife, 10111). After injection, embryos were cultured to the 2-cell stage (24 h) and transferred into pseudopregnant ICR females (8–10 embryos per oviduct). Litters were delivered naturally. Offspring were monitored for growth, health, endocrine status, fertility, and overall development.

### Behavioral assays and sexual activity

Behavioral assays were performed during the light phase after a 1-week habituation.

Open field test^22^: Mice were placed in a 40 × 40 × 40 cm Plexiglass arena for 10 min under 650 lux illumination. Activity was video recorded and analyzed using EthoVision XT. Measures: distance traveled, average speed, and time spent in center vs. periphery. Arenas were cleaned with 70% ethanol between trials.

Novel object recognition test^23^: Day 1: habituation. Day 2: exposure to two identical objects for 10 min. Day 3: one object replaced with a novel one. Recognition index = time with novel / (time novel + time familiar) × 100%.

Object location memory test: Same as NOR, except one object was moved to a new position on Day 3. Location index = time with moved / (time moved + time stationary).

Sexual activity assay^24^: Female mice (≥6 weeks old) were bilaterally ovariectomized under isoflurane anesthesia, given postoperative antibiotics and analgesics, and housed singly until recovery. To induce estrus, females received estradiol benzoate (20 µg in 0.1 mL olive oil, i.p.) 48 h before testing and progesterone (500 µg in 0.1 mL oil, i.p.) 4 h before testing. Estrous status was confirmed by vaginal smears. Sexual behavior tests were conducted in a 40 × 40 × 40 cm Plexiglass open-field arena under dim light at the beginning of the dark cycle. Males were habituated to the apparatus for 2 days.

Interactions with primed females were video-recorded for 30 min. Metrics: mounting latency, intromission latency, ejaculation latency, post-ejaculatory interval, copulatory series number and duration, number of mounts and intromissions. Apparatus was cleaned with 75% ethanol after each test. Observers were blinded to groups.

### Gene expression (qRT-PCR) & Protein analysis

qRT-PCR: RNA was extracted from flash-frozen testes or other tissues with TRIzol (Thermo Fisher 10296028), DNase treated (Turbo DNase, Thermo AM2239), quantified, and reverse transcribed with HiScript III RT SuperMix (Vazyme R323-01). qPCR was performed on a BIO-RAD CFX-96 with SYBR Green master mix (Vazyme Q712-02). Primers spanned exon–exon junctions (see Extended Data Table 1). *Actb* served as housekeeping control. Reactions were run in triplicate. Relative expression was calculated using the ΔΔCq method with efficiency correction^25^. Specificity was validated by melt curves and gel electrophoresis.

Western blot: Proteins were extracted in NP40 buffer (Sangon Biotech 9016-45-9) with protease inhibitors, quantified by BCA assay, separated by SDS–PAGE, transferred to PVDF membranes (PALL BSP0161), and probed with anti-HA (CST C29F4, 1:1000) and anti-GAPDH (Proteintech 60004-1-Ig, 1:5000). HRP-conjugated secondaries and ECL detection (Thermo 34577) were used. Each experiment included 3 biological replicates.

### Hematology and serum biochemistry

Blood was collected via retro-orbital sinus or cardiac puncture. CBCs were measured with Sysmex xn9000. Serum biochemistry (ALT, AST, ALP, bilirubin, BUN, creatinine, electrolytes, glucose) was measured using HITACHI LABOSPECT 008 AS. Analyses were performed blinded to group identity.

### Imaging of LNPs by Transmission Electron Microscopy (TEM)

For TEM observation, 5 μL of each LNP sample was applied to a carbon-coated copper grid and allowed to adsorb for ∼1 min at room temperature. Excess liquid was then removed with filter paper, followed by negative staining with 2% uranyl acetate for 45–60 s. The grid was blotted dry and air-dried before imaging. TEM images were acquired using a Tecnai G2 Spirit BioTwin transmission electron microscope (FEI, USA) operated at 120 kV. Additionally, for fluorescence microscopy assessment, 5 μL of LNP dispersion (with fluorescent tracer incorporated) was spotted on a glass slide, air-dried in the dark for ∼30 min, and then imaged by epifluorescence microscopy to qualitatively examine LNP fluorescence signals.

### Culture and Treatment of Human Seminiferous Tubules

Human seminiferous tubule fragments were obtained from donor testicular tissue under an approved human ethics protocol (No. 20130009 and K2026019) from Zhejiang University human ethics review board. For organ culture, an agarose gel base was prepared by dissolving 0.05 g of agarose in 5 mL ultrapure water, melting it via microwave, and dispensing the solution into a white 96-well plate; the agarose was allowed to cool and solidify at room temperature, forming a half-volume gel layer at the bottom of each well. This soft agarose layer served as a support to maintain an air–medium interface for the explanted tissue^26–28^. Each well was then overlaid with 100 μL of complete culture medium (Procell, CM-H191). Small fragments of human seminiferous tubules were placed in the medium (on top of the gel) and incubated at 33 °C in a humidified 5% CO₂ atmosphere for 1–2 h to equilibrate. Subsequently, Luci MC3 LNP (containing the codon-optimized luciferase mRNA) was added to the wells (typically in a 1:100–1:500 dilution of the stock into the culture medium), and the samples were further incubated under the same conditions for either 24 h or 48 h. After the incubation period (at the designated 24 h or 48 h endpoint), the medium was removed from each well. Then, 100 μL of a detection working solution – composed of 50 μL culture medium mixed with 50 μL of luciferase substrate reagent (Beyotime, Cat. RG043M) – was added to each well, directly covering the tubule fragments. After a 5 min incubation at room temperature to allow substrate penetration and signal development, bioluminescence from the luciferase-expressing tubule fragments was measured using a Spark multimode microplate reader (Tecan), with an integration time of 0.5 s per well. (Luminescence readings were performed for both 24 h and 48 h treated samples to assess transfection efficacy over time.)

### Fluorescence “Squash” Immunostaining

To visualize luciferase expression within the seminiferous tubule cells, a squash immunostaining technique was employed. Cultured human tubule fragments were first fixed in 2% paraformaldehyde (in 1× PBS containing 0.1% Triton X-100) for 5–10 min at room temperature. Each fragment was then gently placed on a glass microscope slide, overlaid with a coverslip, and lightly pressed to flatten the tubule (“squash” preparation). The slide with coverslip was quickly plunged into liquid nitrogen, freezing the sample. Once frozen, the coverslip was pried off, leaving the flattened tubule tissue adhered to the slide. The slides were stored at −80 °C until immunostaining. For staining, slides were brought to room temperature and washed three times with PBS (5 min each). The tissue was then incubated overnight at 4 °C in a humidified chamber with primary antibody against firefly luciferase (rabbit polyclonal, Abcam ab185924, 1:500 dilution in PBS). The next day, slides were washed in PBS three times (10 min each). A secondary antibody (goat anti-rabbit IgG conjugated to Alexa Fluor 488, Invitrogen A11008) diluted 1:500 in PBS was applied to the samples, and slides were incubated for 1–2 h at room temperature in the dark. After secondary incubation, slides were washed again in PBS (3 × 10 min). Finally, samples were mounted with Vectashield antifade medium containing DAPI (Vector Laboratories H-1200) to counterstain nuclei, and a coverslip was applied and sealed with nail polish. Immunofluorescence images were captured using a fluorescence microscope. The resulting images allowed identification of Luci mRNA-translated protein (luciferase) within the tubule cells (green fluorescence), alongside nuclear staining (blue).

### TUNEL Assay for Apoptosis Detection in Testis

Apoptotic cells in mouse testes were detected using a one-step TUNEL (TdT-mediated dUTP nick-end labeling) in situ apoptosis detection kit (Elabscience, Cat. E-CK-A321, FITC Green). Whole testes from mice were fixed in 4% paraformaldehyde (pH 7.4) at 4 °C overnight, then cryoprotected by soaking in 30% sucrose, embedded in OCT compound, and cryosectioned at 5 μm thickness. Frozen sections were equilibrated to room temperature, then post-fixed in 4% PFA for 30 min and washed with PBS (3 × 5 min). Sections were permeabilized with 100 μL of Proteinase K working solution at 37 °C for 20 min, followed by three PBS washes. Next, 100 μL of TdT Equilibration Buffer was applied for 20 min at 37 °C, and subsequently 50 μL of TUNEL reaction mixture (label and enzyme) was added to each section. Slides were incubated in a humidified chamber at 37 °C for 1 h in the dark. For negative-control sections, the TdT enzyme was omitted (to confirm specificity of staining), while for positive controls, sections were pre-treated with DNase I to fragment DNA before TUNEL labeling. After the 1 h incubation, sections were washed with PBS (3 × 5 min), then counterstained with DAPI for 5 min to label nuclei. After four final rinses in PBS, sections were mounted with antifade medium and coverslipped. Fluorescent images of the sections were acquired (TUNEL-positive nuclei appearing as green punctate nuclear fluorescence). Using ImageJ software, the apoptotic index in each section was quantified as described^29^. TUNEL fluorescence intensity (gray value) was measured to indicate the apoptotic signal, while DAPI-stained nuclear area was used to normalize for total cell number. The apoptotic index was calculated as the ratio of TUNEL gray value to total DAPI-positive nuclear area, representing the mean optical density of the TUNEL signal per seminiferous tubule.

### Hormone Measurement by ELISA

Serum hormone levels were measured by enzyme-linked immunosorbent assay (ELISA). In particular, testosterone, luteinizing hormone (LH), and follicle-stimulating hormone (FSH) concentrations in culture media (or serum, as applicable) were quantified using commercial ELISA kits: Testosterone ELISA Kit (SIMI BIO, Cat. ME-M1435S), Luteinizing hormone ELISA Kit (SIMI BIO, Cat. ME-M1128), and FSH ELISA Kit (SIMI BIO, Cat. ME-M1539S). All procedures were carried out according to the manufacturers’ instructions. At the conclusion of the culture or treatment period, samples were collected and incubated with the kit’s assay reagents in 96-well plate format. Absorbance readings were taken on a microplate spectrophotometer, and hormone concentrations were calculated based on the standard curves provided in the kit manuals. Results were reported in ng/mL for testosterone and mIU/mL for FSH, consistent with kit specifications.

### Primers and antibodies

Primer sequences (e.g., *Papolb* forward 5′-CTGTCCATCGACCTGACTTATG-3′; reverse 5′-GCAACTGGTGCAGTTCTTTC-3′; *Actb* forward 5′-TCTAGACTTCGAGCAGGAGATG-3′; reverse 5′-GAACCGCTCGTTGCCAATA-3′) are listed in Extended Data Table 1. Antibody information (host, catalog number, lot, dilution) is also in Extended Data Table 1. Specificity was validated by omission and isotype controls.

### Alkaline Comet Assay

Sperm DNA integrity was assessed using a modified alkaline comet assay (C2041M, Beyotime) adapted for protamine-condensed chromatin. Briefly, sperm washed in Ca²⁺/Mg²⁺-free PBS were embedded in 0.7% low-melting-point agarose on frosted slides. Positive controls were generated by treating sperm with 100 μM H₂O₂ for 30 min at 4 °C. Lysis was performed in two steps. Slides were first incubated in standard lysis buffer (2.5 M NaCl, 100 mM EDTA, 10 mM Tris, 1% Triton X-100, pH 10) for 1 h at 4 °C. This was followed by enzymatic de-protamination in lysis buffer supplemented with 10 mM DTT and 1 mg/mL Proteinase K for 1.5 h at 37 °C. Slides were then equilibrated in alkaline electrophoresis buffer (300 mM NaOH, 1 mM EDTA, pH >13) for 20 min and electrophoresed at 25 V for 10–20 min. After neutralization, DNA was stained with propidium iodide (PI). Images were captured using a fluorescence microscope (Zeiss Axio Imager.M2)

### Masson Trichrome Staining

Masson’s trichrome staining (C0189M, Beyotime) was used to assess tissue fibrosis and collagen deposition. Tissue samples were fixed in 4% paraformaldehyde overnight, dehydrated through a graded ethanol series, and embedded in paraffin. Sections (3 μm thickness) were deparaffinized in xylene and rehydrated using standard protocols. Sections were stained with hematoxylin for 5 min, differentiated in acid alcohol, and blued in running tap water. Slides were then stained with Ponceau–acid fuchsin for 10 min to visualize cytoplasm and muscle fibers (red), followed by differentiation with phosphomolybdic acid. Collagen fibers were counterstained with Light Green solution for 1 min and briefly differentiated in acetic acid. Finally, sections were dehydrated in ethanol, cleared in xylene, and mounted with neutral resin. Slides were evaluated by an investigator blinded to experimental groups. Fibrosis was assessed based on collagen deposition and tissue structural integrity.

### Diff-Quik Staining

Cellular and histological specimens were stained using a Diff-Quik Stain Kit (40748ES, Beyotime) according to the manufacturer’s instructions. For tissue sections, slides were hydrated in distilled water and sequentially immersed in Stain Solution I for 10–20 s and Stain Solution II for 10–20 s with gentle agitation. Slides were rinsed in distilled water and dehydrated twice in absolute ethanol. For smear samples (e.g., sperm smears), slides were air-dried for at least 15 min and fixed in Fixative Solution for 10–20 s. Slides were then stained in Stain Solution I for 5–10 s followed by Stain Solution II for 5–10 s. After rinsing with distilled water and dehydration in ethanol, slides were air-dried, cleared, and mounted for microscopic examination.

### Measurement of mitochondrial membrane potential (Δψm) using JC-1

Mitochondrial membrane potential (Δψm) of spermatozoa was assessed using the JC-1 fluorescent probe (C2003, Beyotime). Sperm concentration was adjusted to 1 × 10⁶ cells/mL and incubated with 1× JC-1 working solution at 37 °C for 20 min in the dark. After staining, cells were washed once with JC-1 staining buffer (500 × g, 5 min), resuspended, and mounted for fluorescence imaging. Green fluorescence (JC-1 monomers, λex/λem ≈ 490/530 nm) and red fluorescence (JC-1 aggregates, λex/λem ≈ 525/590 nm) were captured under identical imaging parameters. Mitochondrial membrane potential was quantified as the ratio of red to green fluorescence intensity. As a positive control for mitochondrial depolarization, samples were pretreated with 10 μM CCCP at 37 °C for 10 min prior to staining.

### TMRE staining for mitochondrial membrane potential

To evaluate Δψm of spermatozoa, cells were stained with tetramethylrhodamine ethyl ester (TMRE; HY-D0985A, MCE). Sperm samples were adjusted to 1 × 10⁶ cells/mL and incubated with 50 nM TMRE for 20 min at 37 °C in the dark. After incubation, cells were washed once with PBS, resuspended, and mounted on glass slides for live-cell fluorescence imaging. All images were acquired using identical excitation wavelength, exposure time, and detector gain to ensure comparability between groups.

### MitoTracker staining

To visualize mitochondrial architecture in spermatozoa, cells were stained with the potential-dependent fluorescent probe MitoTracker Deep Red FM (M22426, Invitrogen). Sperm samples were adjusted to 1 × 10⁶ cells/mL and incubated with 100 nM MitoTracker Deep Red FM for 30 min at 37 °C in the dark. After staining, cells were washed once with PBS (500 × g, 5 min) and mounted for fluorescence microscopy. Mitochondrial signals were captured using a far-red filter set (λex/λem ≈ 644/665 nm). Fluorescence distribution along the sperm midpiece was used to assess mitochondrial integrity and localization.

### Chromosome spread immunostaining

Chromosome spread immunostaining was performed as previously described^30,31^. Briefly, testes were rinsed in cold PBS and the tunica albuginea was removed. Seminiferous tubules were dissociated in DMEM to obtain testicular cell suspensions. Cells were centrifuged at 5800g for 5 min at 4 °C and resuspended in 0.1 M sucrose containing protease inhibitors. Cell suspensions were spread onto slides coated with 1% paraformaldehyde and 0.1% Triton X-100 and incubated overnight in a humid chamber. Slides were washed with PhotoFlo solution and blocked in antibody dilution buffer (ADB). Primary antibodies were applied at 1:500 dilution (mouse anti-SYCP3, ab97672, Abcam; rabbit anti-SYCP1, ab15090, Abcam) and incubated at 37 °C for 2 h. After washing, slides were incubated with Alexa Fluor 488 goat anti-mouse and Alexa Fluor 594 goat anti-rabbit secondary antibodies. Slides were mounted and images were acquired using a Zeiss Axio fluorescence microscope with a 63× oil immersion objective.

### Flow cytometry analysis of testicular cells

To quantify *in vivo* transfection efficiency, single-cell suspensions were prepared from Ai9 mouse testes 48 h after treatment. Decapsulated testes were digested sequentially with collagenase IV (1 mg/mL) and DNase I (0.1 mg/mL) at 33 °C to release interstitial cells, followed by digestion of seminiferous tubules with 0.25% trypsin-EDTA. Cells were filtered through a 70 μm nylon mesh. For germ-cell lineage analysis, cells were stained with Hoechst 33342 (5 μg/mL) for 30 min at 33 °C and propidium iodide (PI) was used to exclude non-viable cells. Data were acquired using a BD LSRFortessa flow cytometer and analyzed with FlowJo software (v10.8). Germ-cell populations were distinguished based on DNA ploidy (1C, 2C, 4C) and cell size (FSC). Transfection efficiency was determined as the percentage of tdTomato-positive cells within each germ-cell population.

### Pyrosequencing

Genomic DNA was extracted using DNeasy Blood and Tissue Kits (69504, Qiagen). Bisulfite conversion was performed using the EpiTect Bisulfite Kit (59104, Qiagen). Primers were designed using PyroMark Assay Design 2.0. Target loci were amplified by PCR and sequencing was performed using the PyroMark Q48 platform following the manufacturer’s instructions with the condition of 95°C for 3 min, followed by 40 cycles of 94°C for 30 s, 56°C for 30 s, and 72°C for 1 min, and a final extension at 72°C for 7 min. Statistical comparisons for individual CpG sites were corrected for multiple testing using the Benjamini–Hochberg method.

### RNA-seq library construction

Total RNA was extracted using the mirVana RNA extraction kit (AM1561, Invitrogen). After DNase treatment (AM2238, Invitrogen), RNA was purified using the RNA Clean & Concentrator-5 Kit (R1013, Zymo Research). Ribosomal RNA was depleted using rDNA probe hybridization followed by RNase H digestion. Strand-specific RNA-seq libraries were prepared using the NEBNext Ultra II Directional RNA Library Prep Kit (E7760S, NEB). Libraries were quantified using Qubit 4.0 and sequenced on a NovaSeq X Plus sequencer (Illumina) with 2 × 150 bp paired-end reads.

### Low-input RNA-seq for early embryo transcriptomics

Two-cell and four-cell embryos were rinsed in PBS and flash-frozen in liquid nitrogen. RNA was extracted using TRIzol reagent followed by column purification using the RNA Clean & Concentrator-5 Kit. DNA contamination was removed using DNase treatment. rRNA depletion and library preparation were performed using the Polaris rRNA/Globin Depletion kit (Watchmaker) and Watchmaker RNA Library Prep Kit. For samples containing <1 ng input RNA, additional bead purification steps were performed using DNA Clean Beads (Vazyme) to remove adapter dimers. Libraries were sequenced on the NovaSeq PE150 platform.

### Whole-genome sequencing

Genomic DNA was fragmented to 180–280 bp using a Covaris instrument. Libraries were constructed using standard Illumina library preparation procedures, including end repair, adaptor ligation, and PCR amplification. Libraries were quality-checked using an Agilent Bioanalyzer 2100 and sequenced on the Illumina NovaSeq platform.

### Bioinformatic analysis

RNA-seq reads were adapter-trimmed using cutadapt^32^. Downstream analyses were performed using piPipes^33^. Transcript abundance was quantified as TPM using the Salmon algorithm^34^, and analyzed with DESeq2 (v.1.46.0)^35^. P-values were adjusted for multiple comparisons using the Benjamini–Hochberg procedure. Differential expression was assessed using adjusted *P* < 0.05. DNA-seq reads were trimmed with cutadapt and aligned to the mouse reference genome (mm10) using Bowtie2^36^. Alignments were converted to BAM format and sorted using SAMtools^37^. Genome-wide read-depth profiles were generated using deepTools^38^. Copy-number variation was assessed by calculating log₂ fold changes in read depth using 1 Mb genomic bins. Structural variants were detected using Picard (Broad Institute; https://broadinstitute.github.io/picard/) and Manta^39^. BAM files were aligned to a hybrid reference genome comprising mm10 and the full *Papolb* cDNA sequence.

### Statistical analysis

All data are shown as mean ± s.d. unless stated otherwise. GraphPad Prism v9 was used. Two-group comparisons: unpaired two-tailed Student’s t-test (if normality passed) or Mann–Whitney U test (if not). Multi-group comparisons: one-way or two-way ANOVA with Tukey’s or Bonferroni post hoc correction. Threshold for significance: *P* < 0.05. Significance reported as *P* < 0.05 (*\*), P < 0.01 (**), P < 0.001 (***)*, and *P < 0.0001 (****)*. Exact *P* values provided where possible. Outliers were tested with Grubbs’ test (α = 0.05). Randomization and blinding were applied where possible. Sample sizes were determined based on previous studies in the field. No animals were excluded except for technical failures, which did not occur in this study.

## Supporting information

Video: MC3 rescued Tes Sperm

## Data availability

The RNA-seq and Genome-seq datasets generated in this study have been deposited in the Genome Sequence Archive (GSA) under BioProject accession PRJCA058762 and Sequence Read Archive accession CRA039361. These data will be made publicly available upon publication.

## Acknowledgments

We thank Yuchao Sun, Ying Liu, Lejun Zhang, Yuting Wang, Wenying Long, Yao Wu, and Yunpan Wang from the Core Facilities of the Fourth Affiliated Hospital, Zhejiang University School of Medicine, and of the International Institutes of Medicine, Zhejiang University, for their technical support. We are grateful to Beijia Wang, Laijie Ni, Xiangfei Zhang, Yulan Zheng, and Weiwei Jin from the Laboratory Animal Center of the International Institutes of Medicine, Zhejiang University, for their meticulous care of the laboratory mice. We also thank the Pathology Core and the Department of Laboratory Medicine of the Fourth Affiliated Hospital, Zhejiang University School of Medicine, as well as members of the Li laboratory, for assistance and helpful discussions. We thank Chenyu Yang in the Center of Cryo-Electron Microscopy (CCEM), Zhejiang University for her technical assistance on Transmission Electron Microscopy. We acknowledge Levostar and GemPharmatech for their contributions to plasmid construction.

## Author Contributions

Qi Jiang and Kexin Su prepared the lipid nanoparticle formulations; Qi Jiang, Yukai Zhong, and Xuliang Ma performed the *in vivo* mouse experiments; Huali Luo, Qi Jiang, Linlin Zhang, and Haifeng Zhang carried out Immunofluorescence Staining, Masson Staining, Diff Quik Staining and H&E Staining; Jiaoyan Ma performed the TUNEL staining and mitochondria related experiment; Qi Jiang, Hao Wang, Qiaodan Li, Yukai Zhong, Jingwen Luo, and Xuliang Ma conducted intracytoplasmic sperm injection and embryo transfer; Qi Jiang and Zhihui Zhang performed the complete blood count and serum biochemistry analyses; Jingwen Luo, Qi Jiang, and Xuliang Ma conducted the behavioral tests; Qi Jiang and Banghua He carried out Western blot assays; Qi Jiang, Yukai Zhong, and Qin Li were responsible for photographing and genotyping the “Papi” and other offspring; Guangnan Li, Wenlin Li, and Chenyu Pei performed RNA-seq and DNA-seq library construction, data analysis, and image processing; Deivendran and Qi Jiang performed Chromosome Spreading; X.Z.L. and Qi Jiang analyzed the data; X.Z.L., Qi Jiang and Guangnan Li wrote the paper; X.Z.L., Xianguang Yang, Lejun Li, Shuai Liu, and Zhefan Yuan contributed to the study design. All authors participated in the preparation of the manuscript.

## Author Information

The authors declare no competing financial interests. Correspondence and requests for materials should be addressed to

