## Extended Data Figure for "Engineering temporal alignment of mRNA delivery enables functional rescue across genetic infertility models"

**This PDF file includes:**

Caption for Extended Data Table 1-4

Extended Data Fig. 1-8

**Other Extended Materials for this manuscript includes the following:**

Extended Data Table 1 (separate file)

Extended Data Table 2 (separate file)

Extended Data Table 3 (separate file)

Extended Data Table 4 (separate file)

Detailed information and statistics for the sequencing data used in this study.

**Extended Data Table 1 (separate file)**

Sexual behavioral parameters of control and F1 (Papi) mice.

**Extended Data Table 2 (separate file)**

**(A)** Primers used in this study for qRT-PCR and genotyping.

**(B)** Antibodies used in the study for western blotting and immunostaining.

**Extended Data Table 3 (separate file)**

**RNA-seq datasets generated in this study.**

**(A)** Differential expression analysis of late two-cell embryos derived from treated versus control sperm.

**(B)** Differential expression analysis of four-cell embryos derived from treated versus control sperm.

**(C)** Correlation analysis of biological replicates for two-cell embryo transcriptomes.

**(D)** Correlation analysis of biological replicates for four-cell embryo transcriptomes.

**(E)** Differential expression analysis of adult heart tissue from F1 offspring derived from treated versus control sires.

**(F)** Differential expression analysis of adult liver tissue from F1 offspring derived from treated versus control sires.

**(G)** Differential expression analysis of adult spleen tissue from F1 offspring derived from treated versus control sires.

**(H)** Differential expression analysis of adult kidney tissue from F1 offspring derived from treated versus control sires.

**(I)** Differential expression analysis of adult testis tissue from F1 offspring derived from treated versus control sires.

Each sheet contains gene identifiers, normalized expression values, $\log_2$ fold changes, raw *P* values, and Benjamini–Hochberg adjusted *P* values (FDR) calculated using DESeq2.

**Extended Data Table 4 (separate file)**

**Whole-genome sequencing–based genomic integrity analysis.**

**(A)** Structural variant detection results based on split-read and discordant-read analysis from whole-genome sequencing of F0 testis samples following mRNA–LNP treatment.

**(B)** Structural variant detection results from whole-genome sequencing of F0 liver samples to assess potential off-target integration events.

**(C)** Structural variant detection results from whole-genome sequencing of F1 testis samples to evaluate potential germline transmission of integration events.

### EXTENDED DATA FIGURE LEGENDS

**Extended Data Figure 1. Inefficient non-viral delivery and context-dependent biodistribution of mRNA in the testis.**

**(a)** Bioluminescence imaging of luciferase reporter expression in adult mouse testes following various non-viral delivery approaches: naked mRNA; mRNA + protamine (pre-condensed); mRNA + electroporation; mRNA + RBCM vesicles; and combinations thereof (including the fully combined method: mRNA + protamine + RBCM + electroporation). Images were acquired at 6, 9, 12, and 24 h after delivery. All treatments remained at background levels comparable to uninjected controls, except the fully combined group, which produced a weak signal. Representative images are shown (n = 3 mice per condition). Dose per testis: 8 µg; injection route: efferent-duct (intratubular) infusion. RBCM, red blood cell membrane.

**(b)** Optimization of protamine:mRNA complex formation by agarose gel-shift assay. Increasing protamine:mRNA mass ratios (w/w) progressively retard free mRNA migration. At a ratio of 1:1 (w/w), no free mRNA band is visible, indicating near-complete complexation. mRNA loaded: 100 ng; gel percentage: 1%.

**(c)** Physicochemical characterization of luciferase mRNA LNPs.
Top: hydrodynamic diameter and polydispersity index (PDI) measured by dynamic light scattering.
Middle: ζ-potential measurements in buffer at pH 7.4, showing near-neutral surface charge (approximately −6 to −3 mV).
Bottom: encapsulation efficiency measured by RiboGreen assay. Data are mean ± s.d. (technical replicates, n = 3). Statistical analysis: one-way ANOVA with Tukey's multiple comparisons test.

**(d)** Transmission electron microscopy (TEM) of LNP formulations. Representative micrographs of MC3-LNP, SM-102-LNP, and ALC-0315-LNP particles show spherical morphology with diameters of ~100 nm. Scale bar, 100 nm. (n = 9 images from 3 independent preparations.)

**(e)** *Ex vivo* organ bioluminescence 12 h after intratubular injection of luciferase mRNA LNPs, shown as photon flux (photons $s^{-1}$ $cm^{-2}$ $sr^{-1}$; logarithmic scale). Data are mean ± s.d. (n = 3 mice per group). Statistical analysis: one-way ANOVA with Tukey's multiple comparisons test.

**(f)** *In vivo* bioluminescence imaging 1–24 h after intratubular injection of luciferase mRNA formulated in SM-102 LNP at four doses (1.25, 2.5, 5, and 10 µg per testis). Luminescence in the liver/spleen region is visible even at the lowest dose (1.25 µg) and increases with dose. Representative images are shown (n = 3 mice per dose).

**(g)** Gross appearance of testes 12 h after intratubular luciferase mRNA LNP injection. The SM-102-treated testis is enlarged and erythematous compared with controls, whereas MC3- and ALC-0315-treated testes show no obvious inflammation. Other organs appear similar across groups. Scale bar, 1 cm. (n = 3 mice per group.)

**(h)** H&E-stained sections of major organs (liver, spleen, kidney, heart, lung, and epididymis) 12 h after intratubular injection of luciferase mRNA LNPs (dose: 8 µg). SM-102-treated mice exhibit mild hepatic sinusoidal dilation, whereas other organs appear normal. Scale bar, 100 µm. (n = 3 mice per group; representative of 3 sections per organ.)

**(i)** Ai9 Cre-reporter schematic (top). *Ex vivo* fluorescence imaging of Ai9 mice 48 h after intratubular injection of Cre mRNA LNPs (bottom). H, heart; Lu, lung; Li, liver; S, spleen; K, kidney; T, testis. Representative images are shown (n = 3 mice per group).

**(j)** *Ex vivo* fluorescence imaging of major organs from Ai9 mice 48 h after intratubular injection of Cre mRNA LNPs (MC3, ALC-0315, SM-102; n = 3 mice per group). SM-102 shows off-target tdTomato expression in liver and spleen.

**(k)** Gross images of harvested major organs at 48 h after injection. Scale bar, 1 cm.

**(l)** H&E-stained sections 48 h after intratubular injection of 8 µg Cre mRNA LNPs. SM-102-treated testes show inflammatory features, whereas MC3- and ALC-0315-treated testes appear normal. Scale bar, 100 µm. (n = 3 mice per group; representative of 3 sections.)

**(m)** Flow-cytometry gating strategy for identification of viable singlets and germ-cell subpopulations based on DNA content (1C, 2C, 4C) and forward scatter.

**(n)** Flow-cytometry quantification of tdTomato expression. Left: mean fluorescence intensity (MFI) of tdTomato in each germ-cell population for each LNP formulation. Right: proportion of cells within each spermatogenic stage. Data are mean ± s.d. (n = 6 testes per group). Statistical analysis: two-way ANOVA with Tukey's multiple comparisons test.

**(o)** Representative seminiferous tubule cross-section co-stained for tdTomato (red), SOX9 (Sertoli-cell marker; green), and nuclei (DAPI, blue). Scale bar, 40 µm. Representative images are shown (n = 3 mice per group; 3 sections per testis).

**(p)** Summary of delivery kinetics and functional outcomes across LNP formulations.

Jiang et al, Extended Data Fig. 1

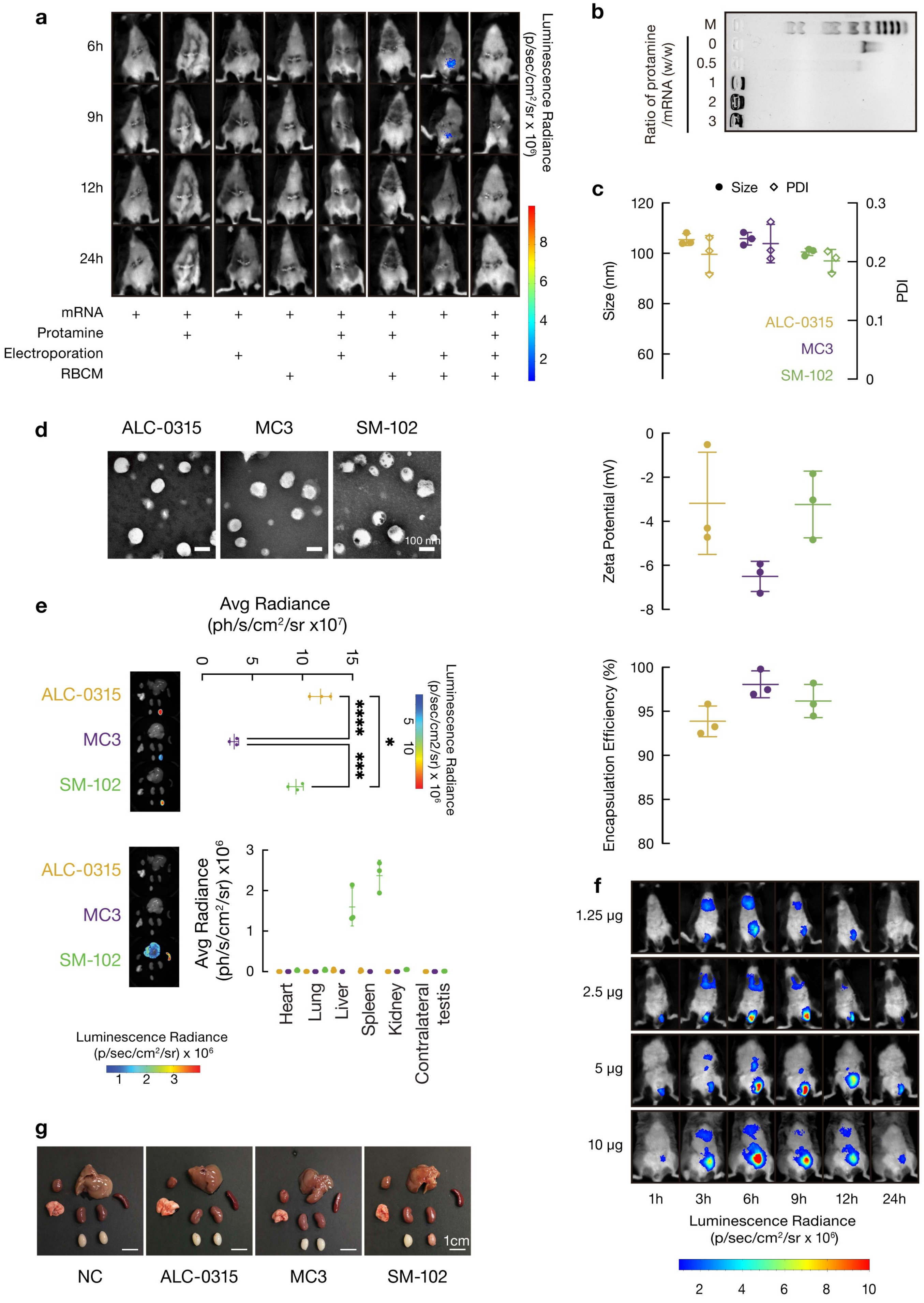

Jiang et al, Extended Data Fig. 1

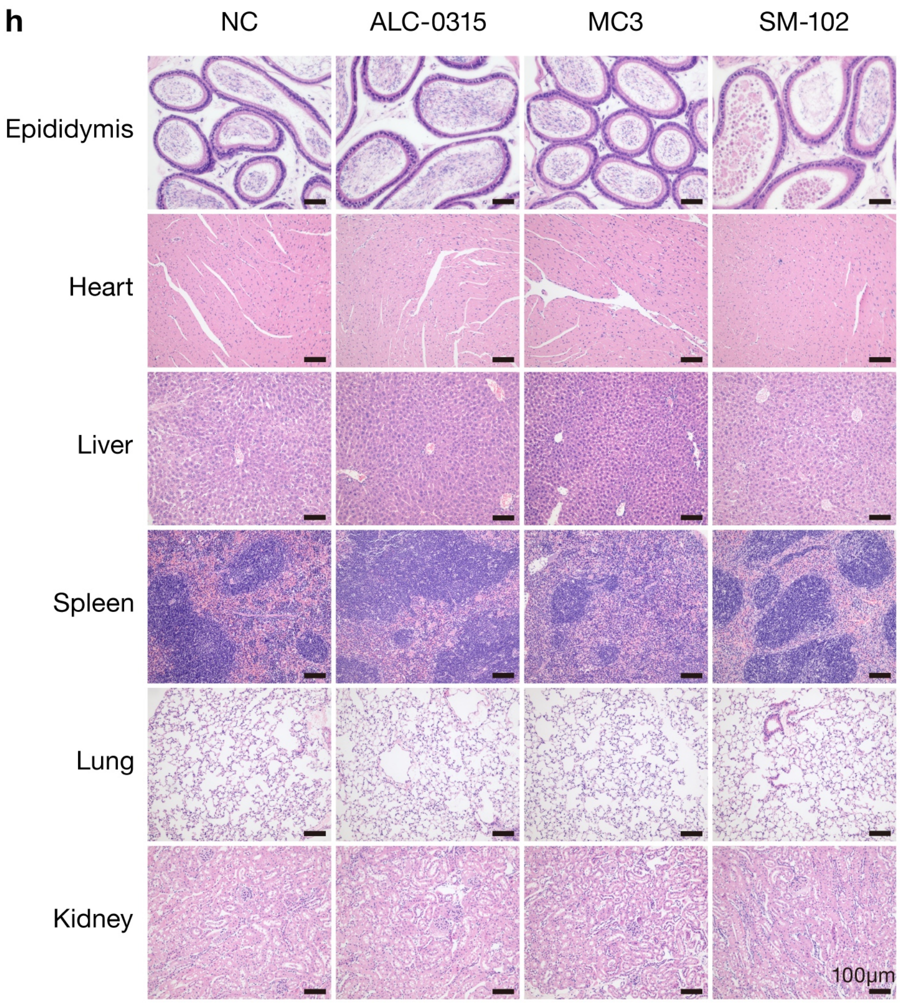


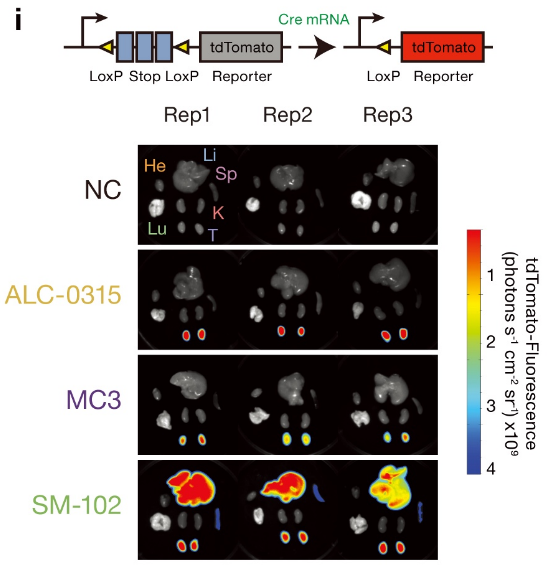

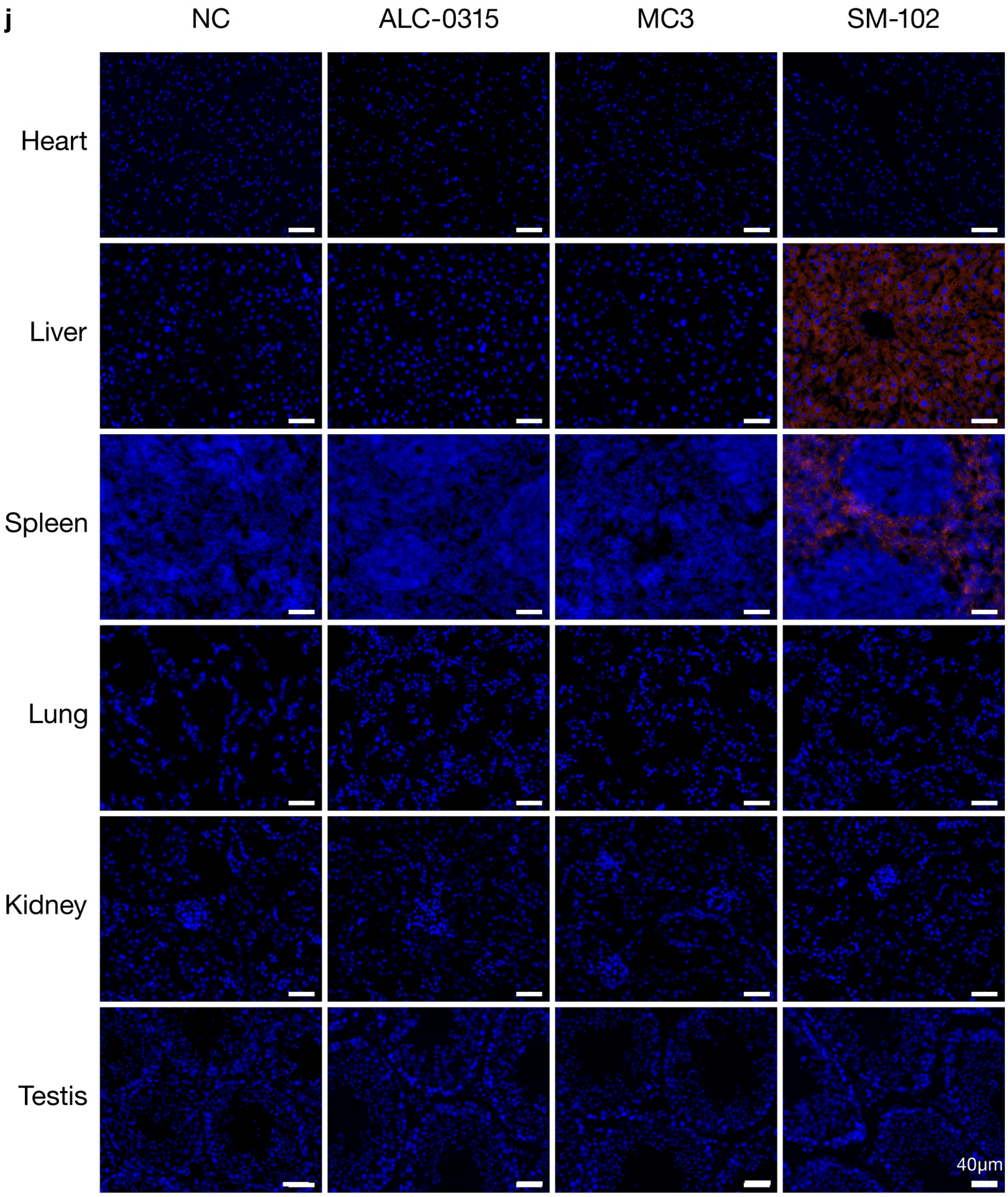
j
NC
ALC-0315
MC3
SM-102
Heart
Liver
Spleen
Lung
Kidney
Testis
40μm

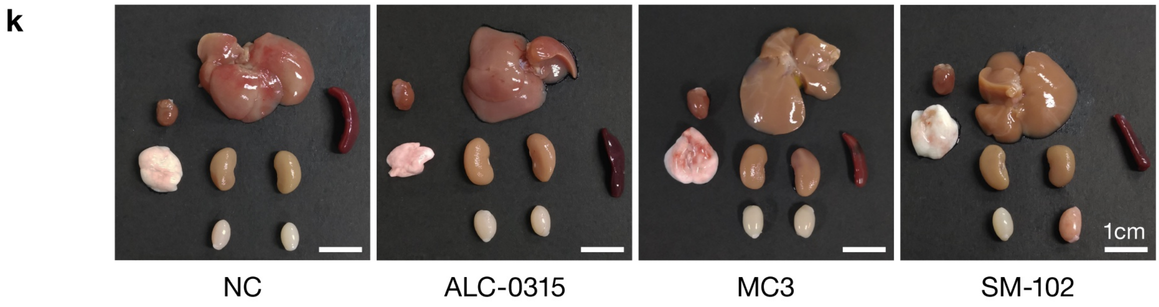
k
1cm
NC
ALC-0315
MC3
SM-102

Jiang et al, Extended Data Fig. 1

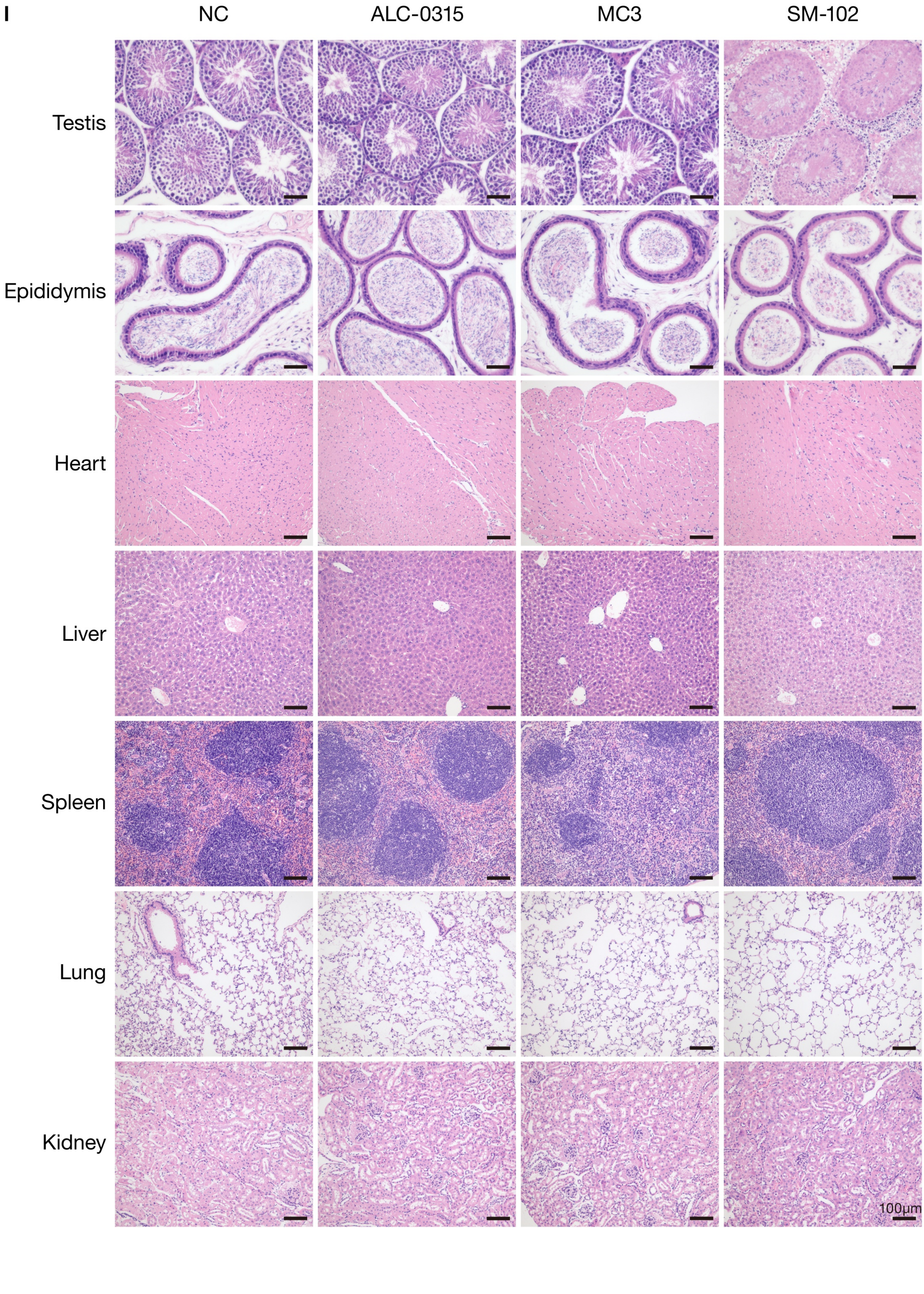

Jiang et al, Extended Data Fig. 1

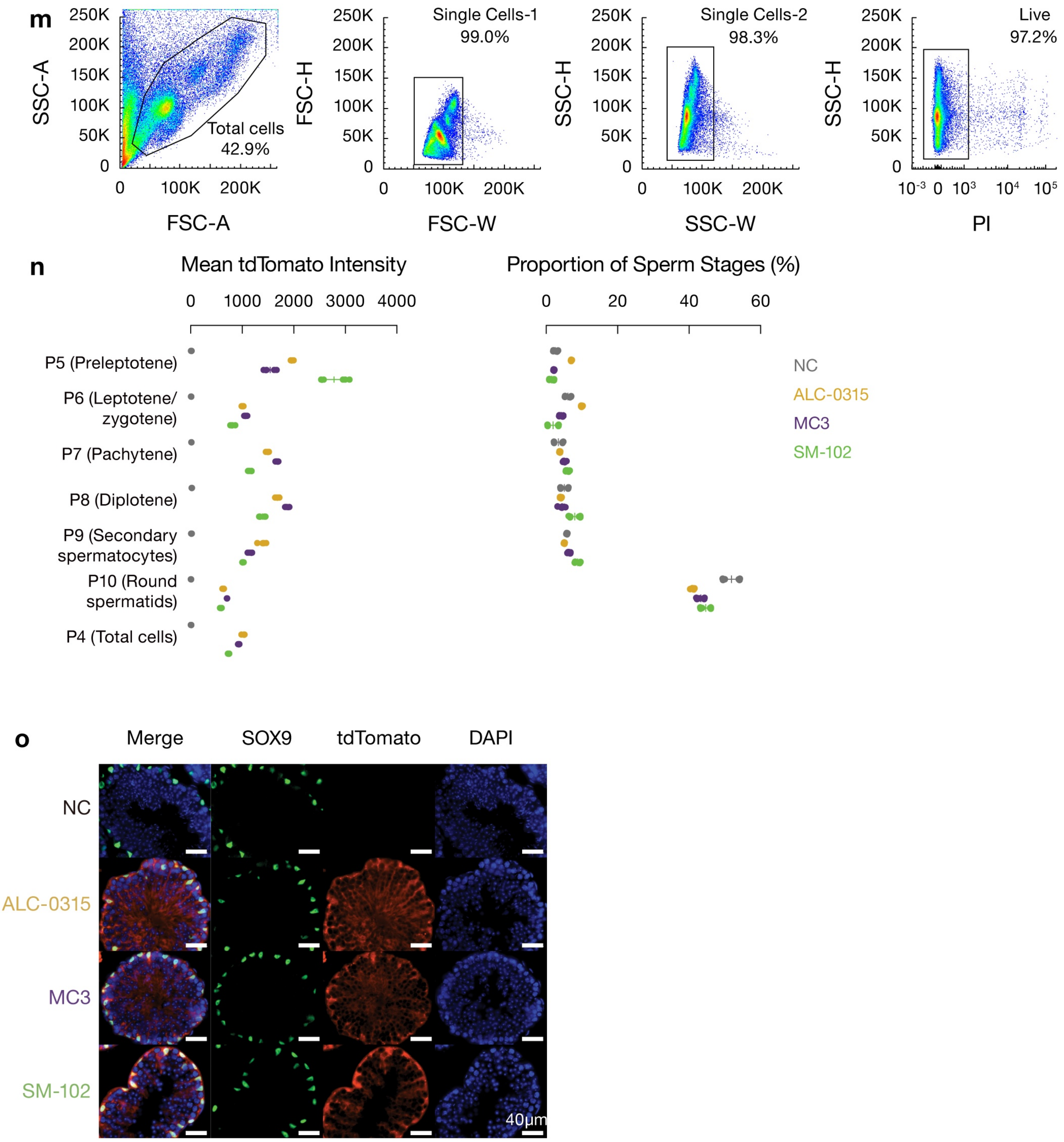


p

| **Parameter** | **MC3** | **ALC-0315** | **SM-102** |
|---|---|---|---|
| Peak expression | Moderate | High | High |
| Expression duration | Sustained (~10–14 d) | Short (~3–5 d) | Short |
| Inflammation | Minimal | Minimal | High |
| Functional outcome | Rescue | None | None |

**Extended Data Figure 2. RNA format and chemical modification influence expression kinetics and immunogenicity *in vivo***

**(a)** Representative fluorescence images illustrating GFP expression kinetics among RNA formats at indicated time points.

**(b)** Quantification of GFP expression kinetics in testes following delivery of different RNA formats (MC3 LNP): $N^1m\Psi$-modified eGFP mRNA, saRNA, and circRNA. Data are mean ± s.d. (n = 3 mice per group). Statistical analysis: two-way ANOVA with Tukey's multiple comparisons test.

**(c)** Innate immune response in the testis 24 h after delivery of different RNA formats (*Il-10*, *Il-12b*, *Tnf-α*, *Cxcl10*, *Il-6*, *Ifnb1*, *Il-1β*; internal reference: *β-actin*). Data are mean ± s.d. (n = 3 mice per group). Statistical analysis: one-way ANOVA with Tukey's multiple comparisons test.

**(d)** Representative fluorescence images comparing unmodified versus $m^1\Psi$-modified eGFP mRNA expression at indicated time points.

**(e)** Time-course quantification of unmodified versus modified eGFP mRNA expression. Data are mean ± s.d. (n = 3 mice per group). Statistical analysis: two-way ANOVA with Tukey's multiple comparisons test.

**(f)** RT–qPCR measurement of eGFP mRNA abundance in the testis at indicated time points after injection of unmodified versus modified eGFP mRNA. Normalization gene: *β-actin*. Data are mean ± s.d. (n = 3 mice per group). Statistical analysis: two-way ANOVA with Tukey's multiple comparisons test.

**(g)** Induction of innate immune genes at 6, 12, and 24 h after injection of unmodified versus modified eGFP mRNA. Data are mean ± s.d. (n = 3 mice per group). Statistical analysis: one-way ANOVA with Tukey's multiple comparisons test.

Jiang et al, Extended Data Fig. 2

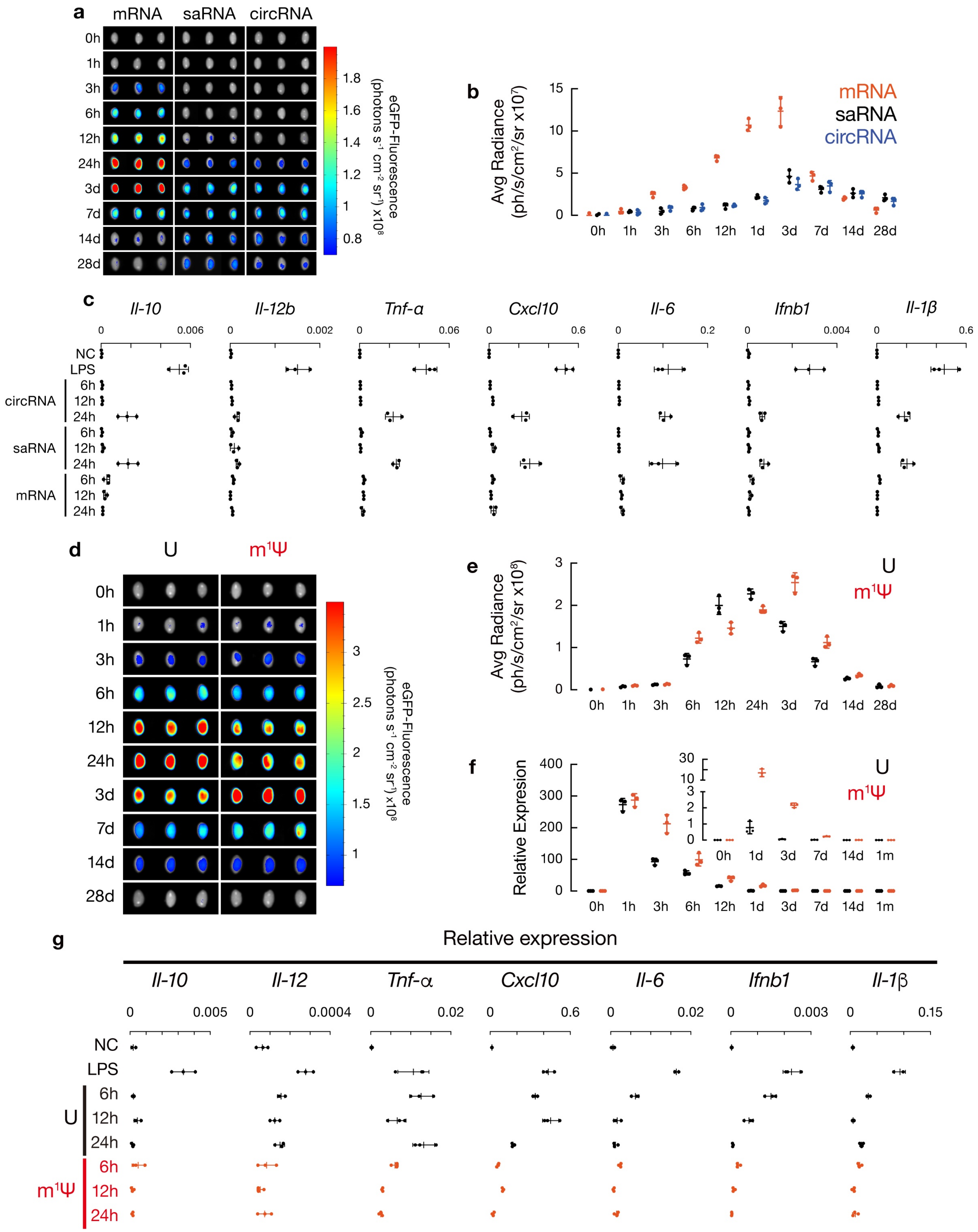

**Extended Data Figure 3. Quality control and tissue-restricted expression of *Papolb* mRNA.**

**(a)** *Papolb* plasmid construct (left), linearized plasmid used for *in vitro* transcription (middle), and quality assessment of the *in vitro*–transcribed *Papolb* mRNA (right). The gel image shows intact full-length transcript with minimal degradation. Representative image from 3 independent transcription reactions.

**(b)** Quality control of *in vitro*–transcribed mRNAs used in this study.
Top: double-stranded RNA (dsRNA) content measured using dsRNA Quantification Kit (Hzymes, HBP003805) (dsRNA <0.005%). dsRNA, double-stranded RNA.
Bottom: endotoxin levels measured by LAL assay (<4 EU $mL^{-1}$). Data represent 3 independent preparations. LAL assay, limulus amebocyte lysate assay.

**(c)** Western blot analysis of HA–PAPOLB expression in indicated organs 10 h after intratubular delivery of MC3–*Papolb* mRNA. HA–PAPOLB protein was detected only in the injected testis but not in contralateral testis or other organs. GAPDH serves as loading control. (n = 3 mice).

**(d)** H&E staining of wild-type testes three weeks after intratubular injection of MC3–*Papolb* or ALC-0315–*Papolb* mRNA LNPs compared with untreated *Papolb*$^{+/-}$ controls. Testicular morphology remains normal under both conditions. Scale bar, 50 µm. (n = 3 mice per group; representative of 3 sections per testis).

Jiang et al, Extended Data Fig. 3

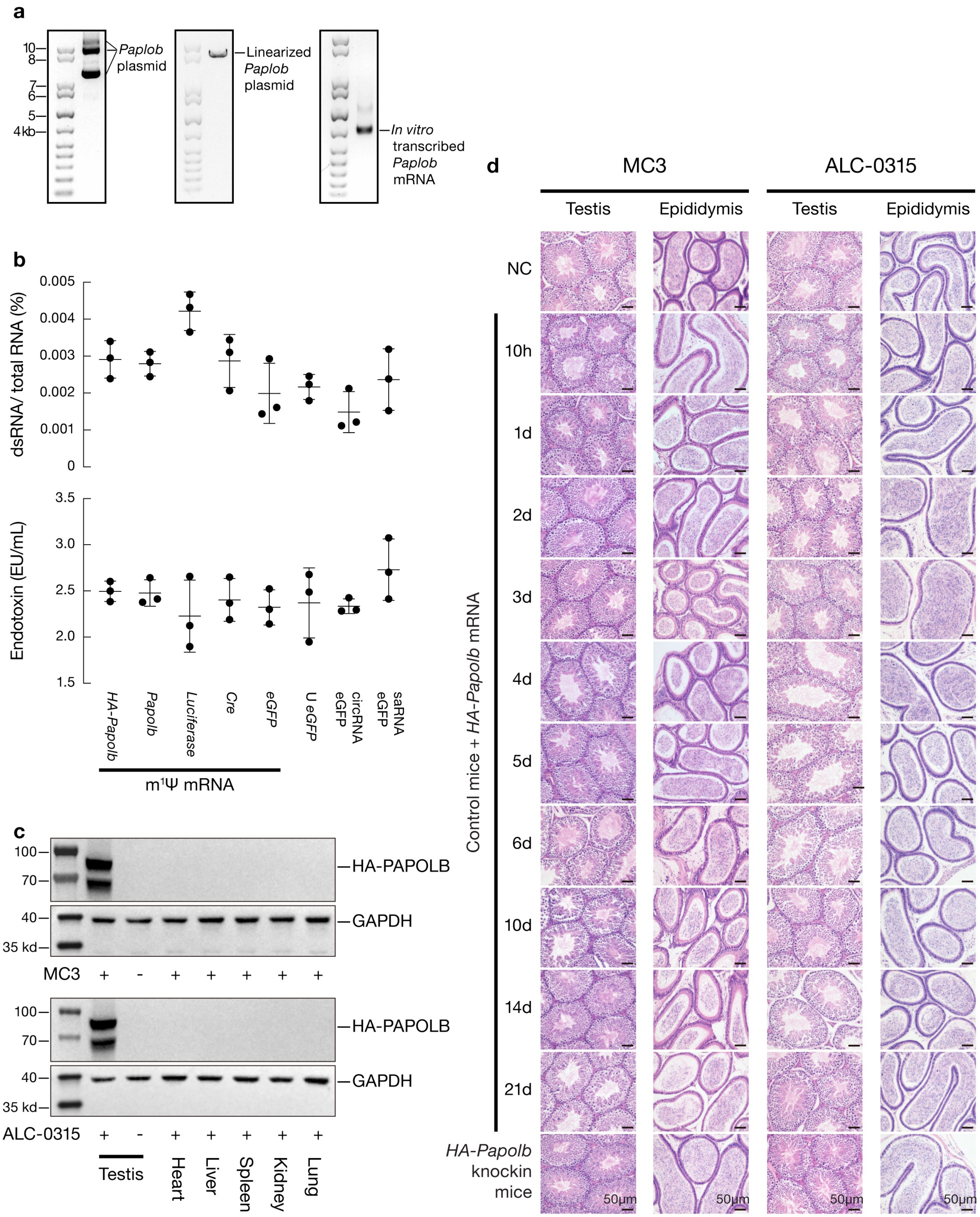

**Extended Data Figure 4. Stage-specific markers and temporal progression of *Papolb*-mediated spermatogenic rescue**

**(a)** H&E-stained testis sections across time points after a single 8 µg ALC-0315–*Papolb* mRNA injection. Scale bar, 40 µm. (n = 3 mice per time point; representative of 3 sections per testis).

**(b–e)** Immunofluorescence markers of spermatogenic progression at 11 days post-treatment in *Papolb*$^{-/-}$ untreated and treated testes compared with *Papolb*$^{+/-}$ controls:
**(b)** TNP1/H3;
**(c)** ACRV1/SYCP3;
**(d)** SYCP1/γH2AX;
**(e)** H3/PCNA.

ACRV1 and TNP1 label elongating spermatids in treated KO testes, whereas untreated KO testes lack these signals. PCNA, SYCP3, SYCP1, and γH2AX show comparable distributions across groups. Scale bar, 40 µm. (n = 3 mice per group; representative of 3 sections per testis).

**(f)** PNA (green) and Protamine-1 (red) staining in testis sections at day 7 after treatment across different dose groups. Scale bar, 20 µm. (n = 3 mice per group; all tubules within each section were analyzed).

**(g)** TUNEL staining of seminiferous tubules following escalating doses of MC3–*Papolb* mRNA (0–16 µg). Scale bar, 100 µm. (n = 3 mice per group; 10 tubules quantified per section). TUNEL, Terminal deoxynucleotidyl transferase dUTP nick end labeling.

**(h)** Innate immune response in testes following delivery of different *Papolb* mRNA doses. Data are mean ± s.d. (n = 3 mice per group). Statistical analysis: one-way ANOVA with Tukey's multiple comparisons test.

**(i)** Time-course quantification following a single 8 µg MC3–*Papolb* mRNA injection: percentage of seminiferous tubules containing elongating spermatids, seminiferous epithelial height, and Johnsen score. Data are mean ± s.d. (n = 3 mice per time point). Statistical analysis: one-way ANOVA with Tukey's multiple comparisons test.

**(j)** Representative PNA/Protamine-1 staining across time points after treatment. Scale bar, 40 µm.

**(k)** Representative high-magnification images illustrating spermatid developmental steps (steps 9–11 at day 7; steps 13–14 at day 9; step 15 at day 11; step 16 at day 13). Scale bar, 40 µm.

**(l)** Histological impact of repeated MC3–LNPs injections in *Papolb*$^{-/-}$ testes. At day 13 and day 17 after repeated intratubular injections, some seminiferous tubules contain elongated sperm in the lumen, whereas others show germ-cell depletion and structural disruption. Scale bars, 40 µm and 100 µm. (n = 3 mice per group).

Jiang et al, Extended Data Fig. 4

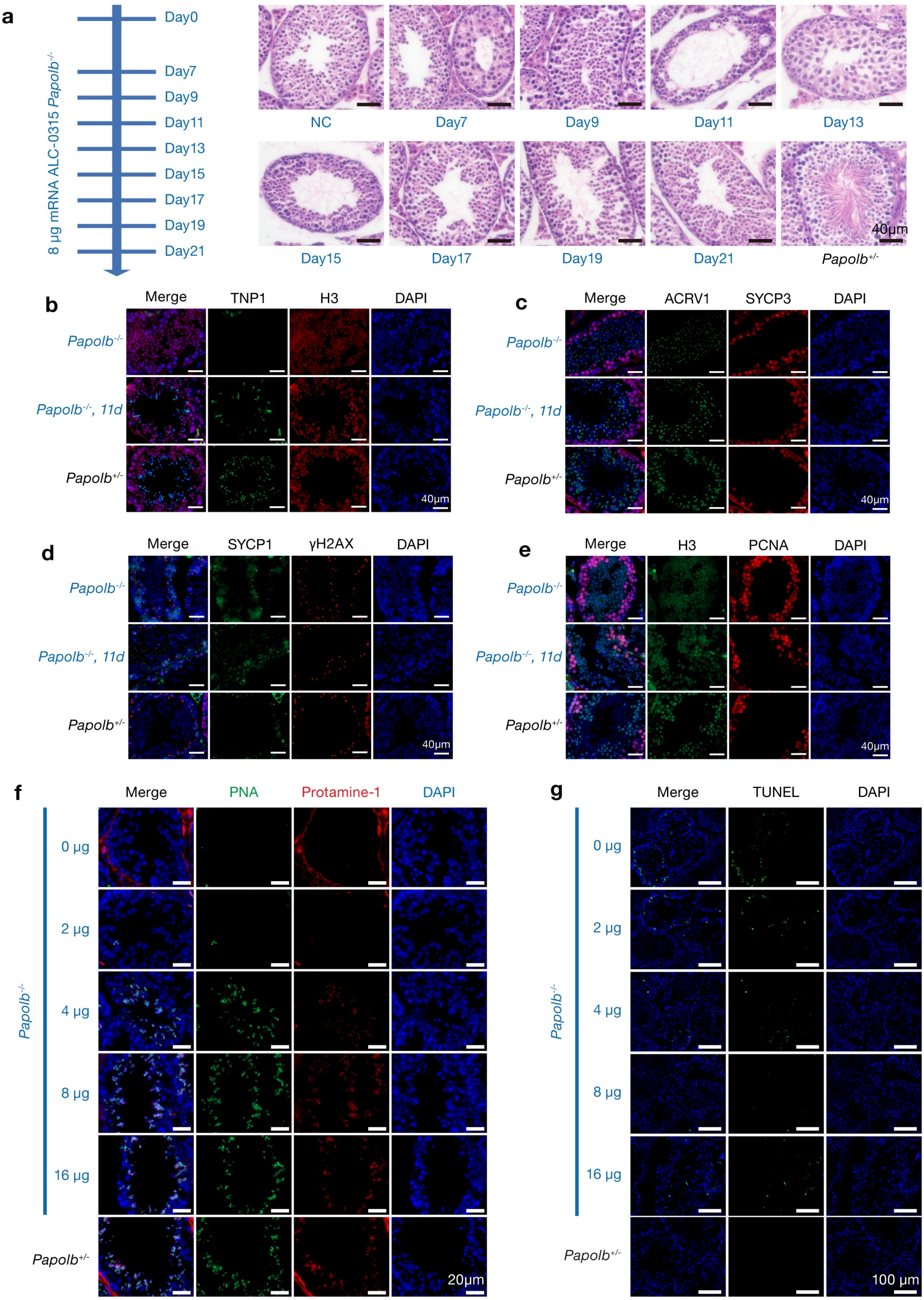

Jiang et al, Extended Data Fig. 4

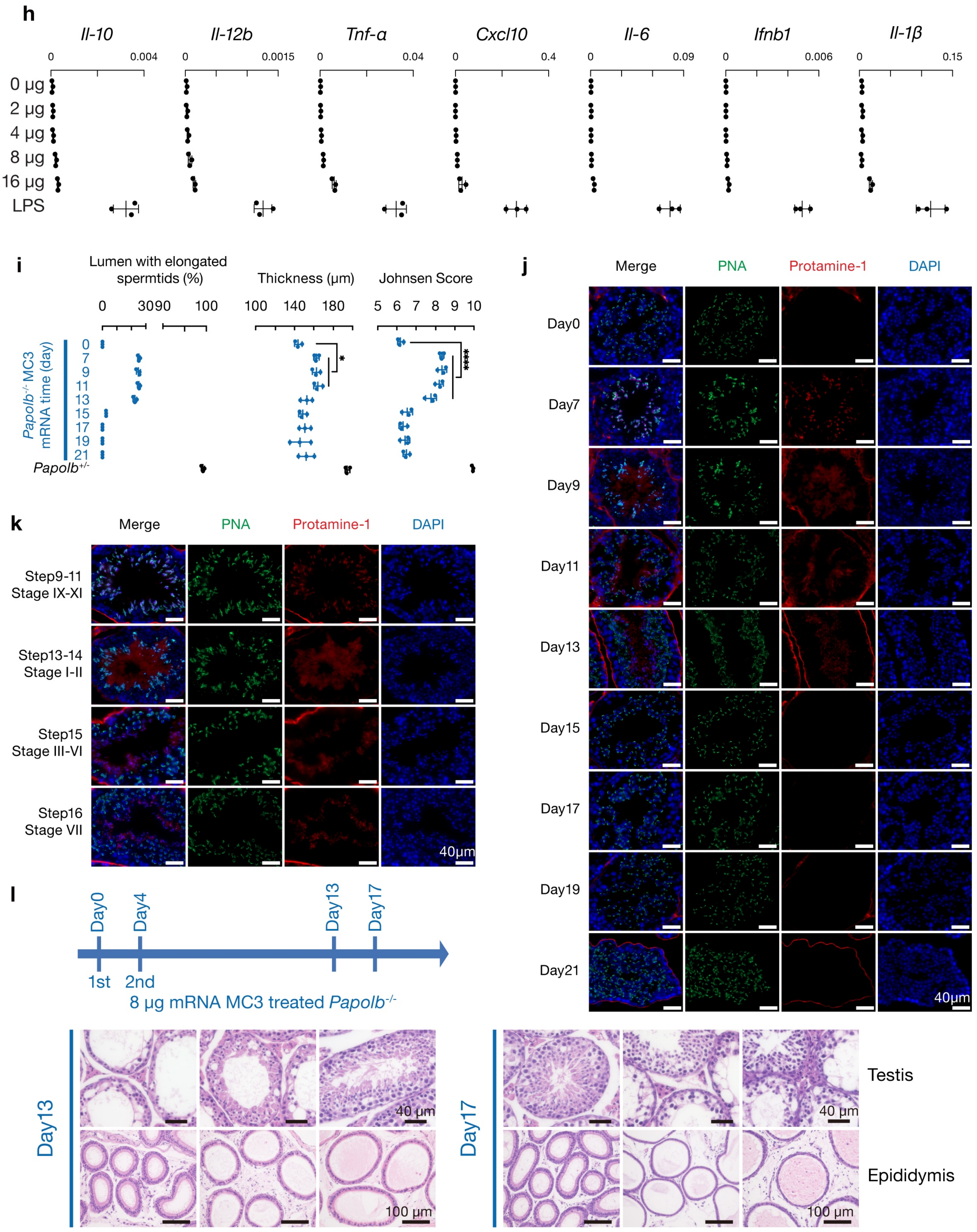

**Extended Data Figure 5. Functional competence of rescued sperm and normal physiological outcomes in offspring.**

**(a–c)** Epididymal histology across dose and time points. No mature sperm were detected in epididymal lumens of treated KO mice, consistent with disruption of sperm transit following intratubular injection. Scale bar, 50 µm. (n = 3 mice per group).

**(d–e)** Mitochondrial assessments in testicular sperm using (a) MitoTracker and (b) TMRE staining. Sperm from treated *Papolb*$^{-/-}$ mice were compared with *Papolb*$^{+/-}$ controls. Scale bar, 10 µm. (n = 3 mice per group; ≥200 sperm analyzed). TMRE, tetramethylrhodamine ethyl ester perchlorate.

**(f)** Embryo development following ICSI using rescued sperm. Representative images of embryos at zygote (Day 0), two-cell (Day 1), cleavage/morula (Days 2–3), and blastocyst (Day 4.5) stages. Example dataset: 40 injected oocytes, 34 two-cell embryos, and 24 blastocysts.

**(g)** Representative offspring generated by ICSI using rescued *Papolb*-deficient sperm at different postnatal ages.

**(h–i)** Gross morphology (scale bar, 2 cm) and H&E staining of major organs from Het and Papi mice. Scale bar, 100 µm. (n = 3 mice per group).

**(j)** DEXA analysis of body composition in Het and Papi F1 mice at 10 weeks of age. Parameters include body weight, fat mass fraction, lean mass fraction, bone mineral density, and bone mass fraction. Data are mean ± s.d. (n = 6 mice per group). Statistical analysis: unpaired t-test.

**(k)** Complete blood count analysis. Parameters include WBC, RBC, HGB, PLT, HCT, MCV, MCH, and MCHC. Data are mean ± s.d. (n = 5 mice per group). No significant differences were observed between Het and Papi groups ($P > 0.05$). WBC, white blood cell count; RBC, red blood cell count; HGB, hemoglobin; PLT, platelet count; HCT, hematocrit; MCV, mean corpuscular volume; MCH, mean corpuscular hemoglobin; MCHC, mean corpuscular hemoglobin concentration.

**(l)** Serum biochemical analysis assessing liver enzymes (ALT, AST, ALP, T-Bil), kidney function (BUN, CRE), Glucose and LDH. Data are mean ± s.d. (n = 5 mice per group). Statistical analysis: unpaired t-test. ALT, alanine aminotransferase; AST, aspartate aminotransferase; ALP, alkaline phosphatase; T-Bil, total bilirubin; BUN, blood urea nitrogen; CRE, creatinine; LDH, lactate dehydrogenase.

**(m)** Behavioral assays comparing Het and Papi F1 mice, including open-field activity, object location memory, and novel object recognition. Data show no significant differences between groups (n = 5 mice per group). Statistical analysis: unpaired t-test.

**(n)** Sex ratio of F1 offspring generated by ICSI using epididymal sperm or testicular sperm from *Papolb*$^{+/-}$ mice, or testicular sperm from mRNA-treated *Papolb*$^{-/-}$ mice. (n = 31, 33, and 29 offspring, respectively; Chi-square test).

**(o–p)** Representative images of F2 and F3 offspring derived from rescued *Papolb* lineages.

**(q)** PCR genotyping of offspring from MC3-treated KO fathers. All offspring carried one mutant and one wild-type allele, confirming expected Mendelian inheritance and absence of transgene integration.

**(r)** Serum testosterone and FSH levels measured by ELISA in Het, Papi F1, and Papi F2 mice. Data are mean ± s.d. (n = 5 mice per group). Statistical analysis: one-way ANOVA with Tukey’s multiple comparisons test. FSH, follicle-stimulating hormone.

Jiang et al, Extended Data Fig. 5

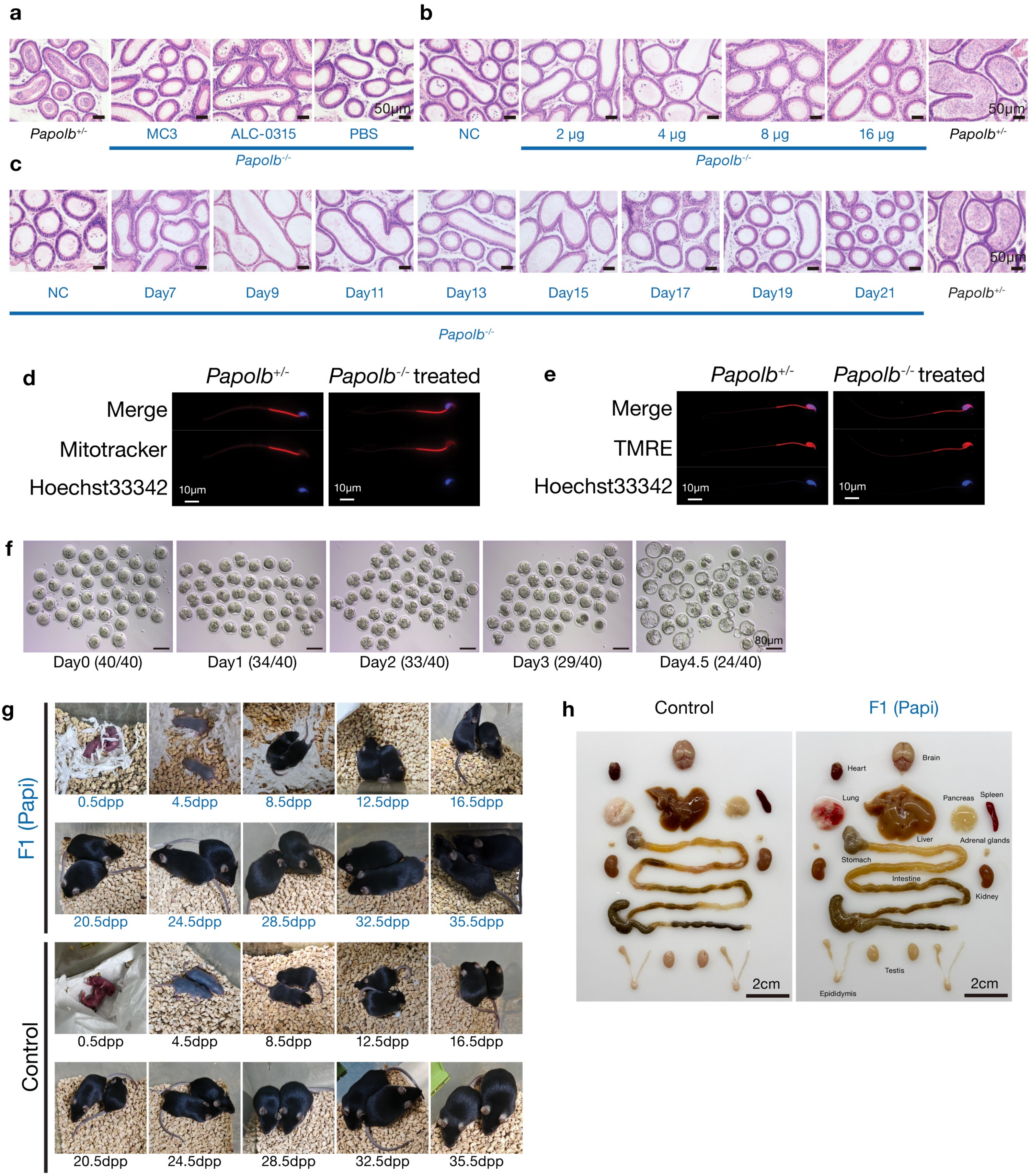

Jiang et al, Extended Data Fig. 5

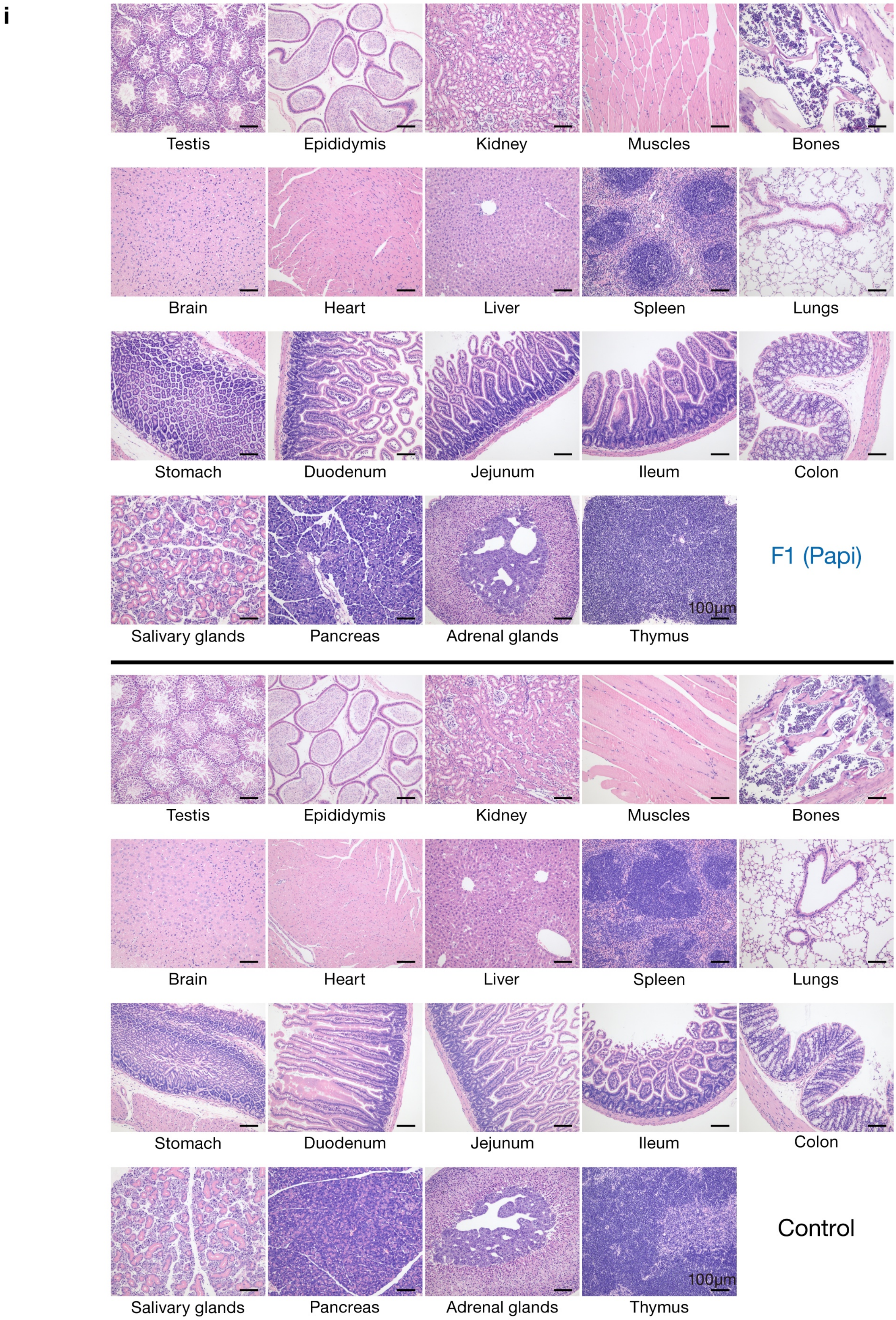

Jiang et al, Extended Data Fig. 5

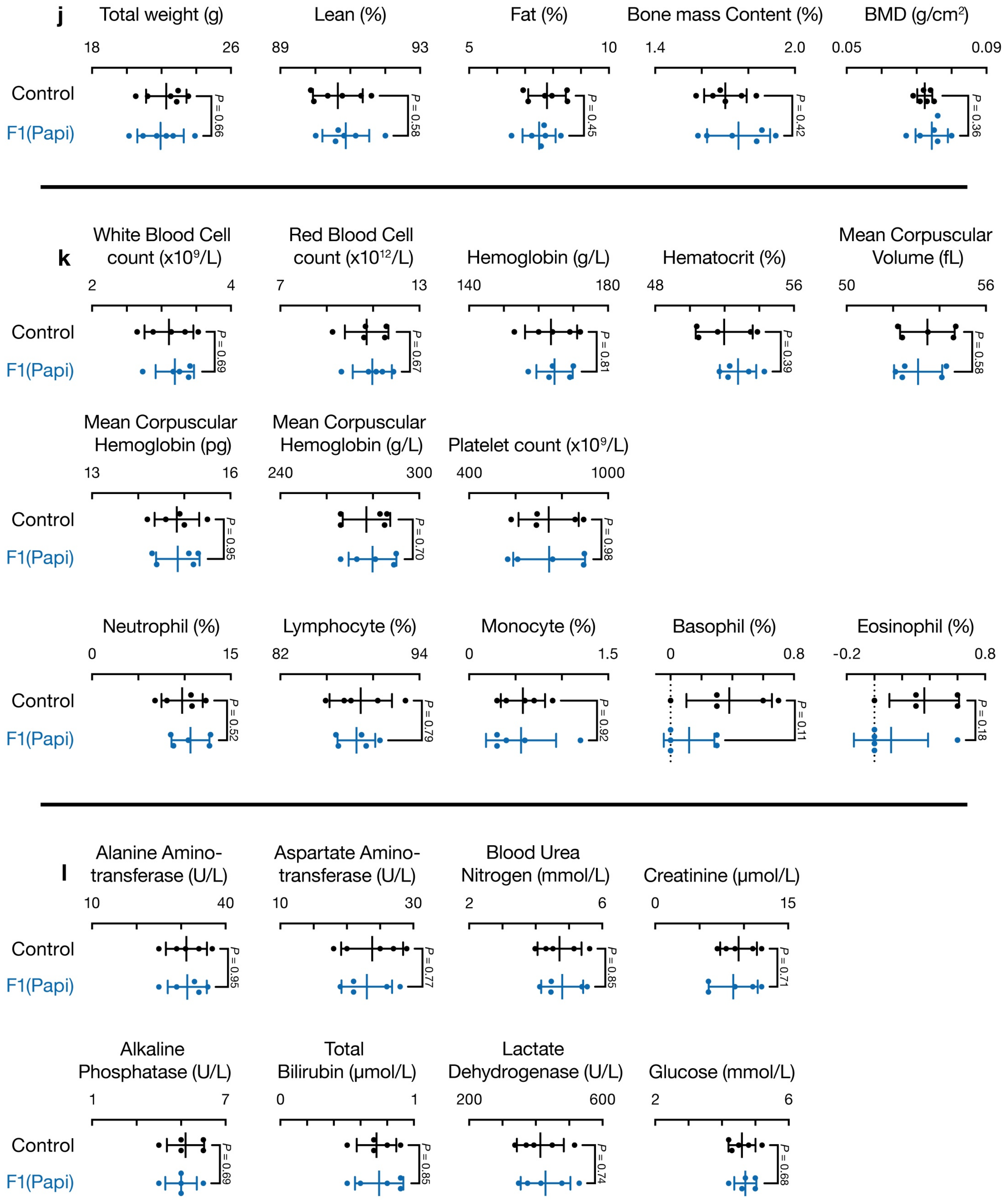

Jiang et al, Extended Data Fig. 5

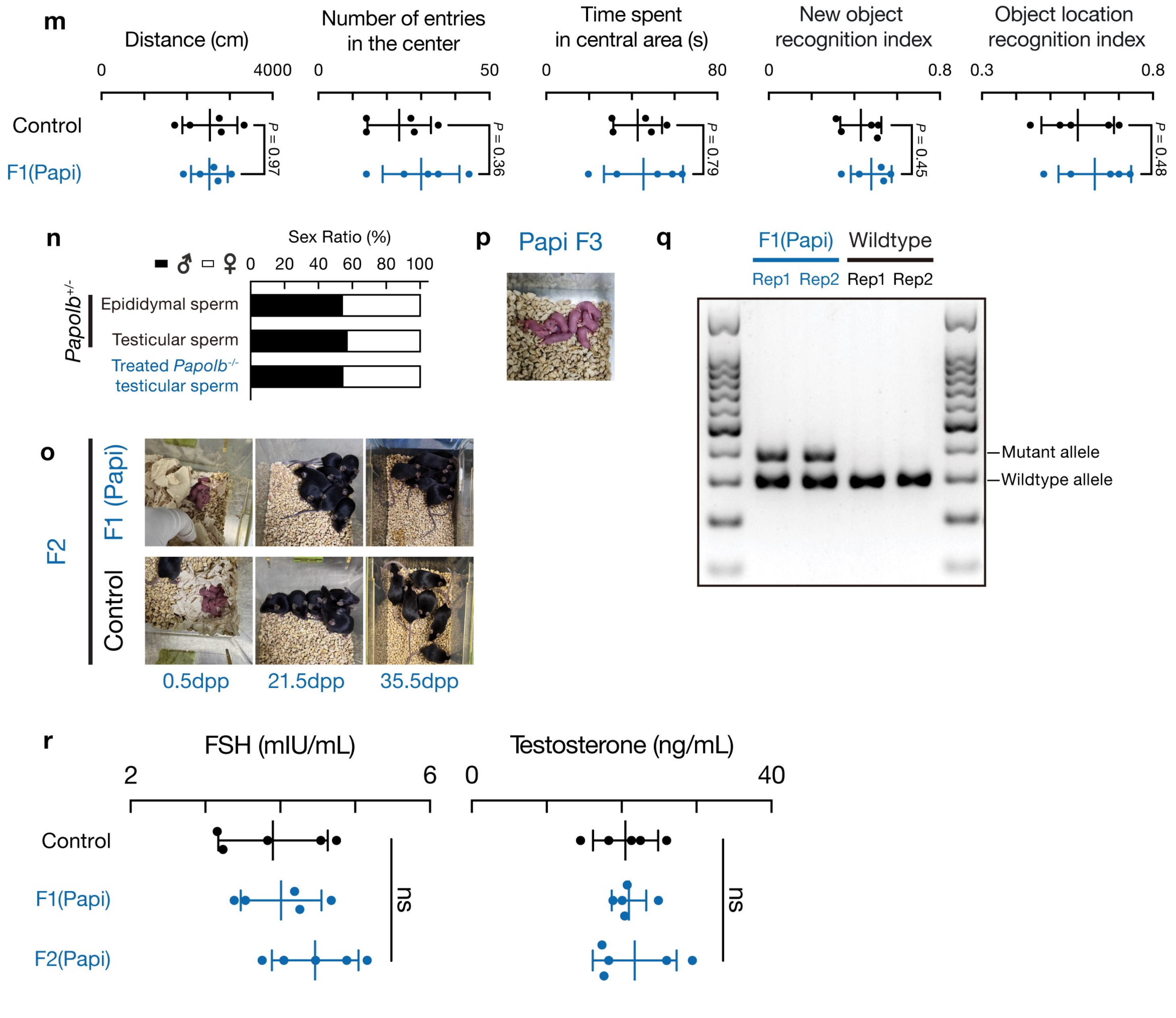

**Extended Data Figure 6. Functional rescue and developmental progression in *Spo11* and *Btbd18* infertility models**

**(a–b)** Time-course immunofluorescence staining for PNA (green) and Protamine-1 (red) in *Spo11*$^{-/-}$ and *Btbd18*$^{-/-}$ rescue experiments. Scale bar, 40 μm. (n = 3 mice per group).

**(c)** Immunofluorescence markers of spermatogenic progression (ACRV1, TNP1, γH2AX, PCNA) following rescue. Scale bar, 40 μm. (n = 3 mice per group).

**(d)** Quantification of rescued tubules (PNA-positive) at peak time points for *Spo11* (left) and *Btbd18* (right) rescue. Tubules counted as 100 tubules per section × 3 sections per testis (n = 3 mice per group). Statistical analysis: one-way ANOVA with Tukey's multiple comparisons test.

**(e)** Epididymal H&E sections at indicated time points showing absence of mature sperm. Scale bar, 40 μm. (n = 3 mice).

**(f–g)** Sperm morphology and two-cell embryos generated via ICSI using sperm from treated *Spo11*$^{-/-}$ and *Btbd18*$^{-/-}$ males. Scale bars, 10 μm and 80 μm. (n = 4 sessions; ~20 embryos per group).

**(h–i)** Representative F1 offspring and PCR genotyping confirming expected heterozygous inheritance.

**(j)** Sex ratio of F1 offspring generated by ICSI using testicular sperm from *Spo11*$^{+/-}$ and *Btbd18*$^{+/-}$ mice, or testicular sperm from mRNA-treated *Spo11*$^{+/-}$ and *Btbd18*$^{+/-}$ mice. (n = 30, 31, 33, and 36 offspring, respectively; Chi-square test).

Jiang et al, Extended Data Fig. 6

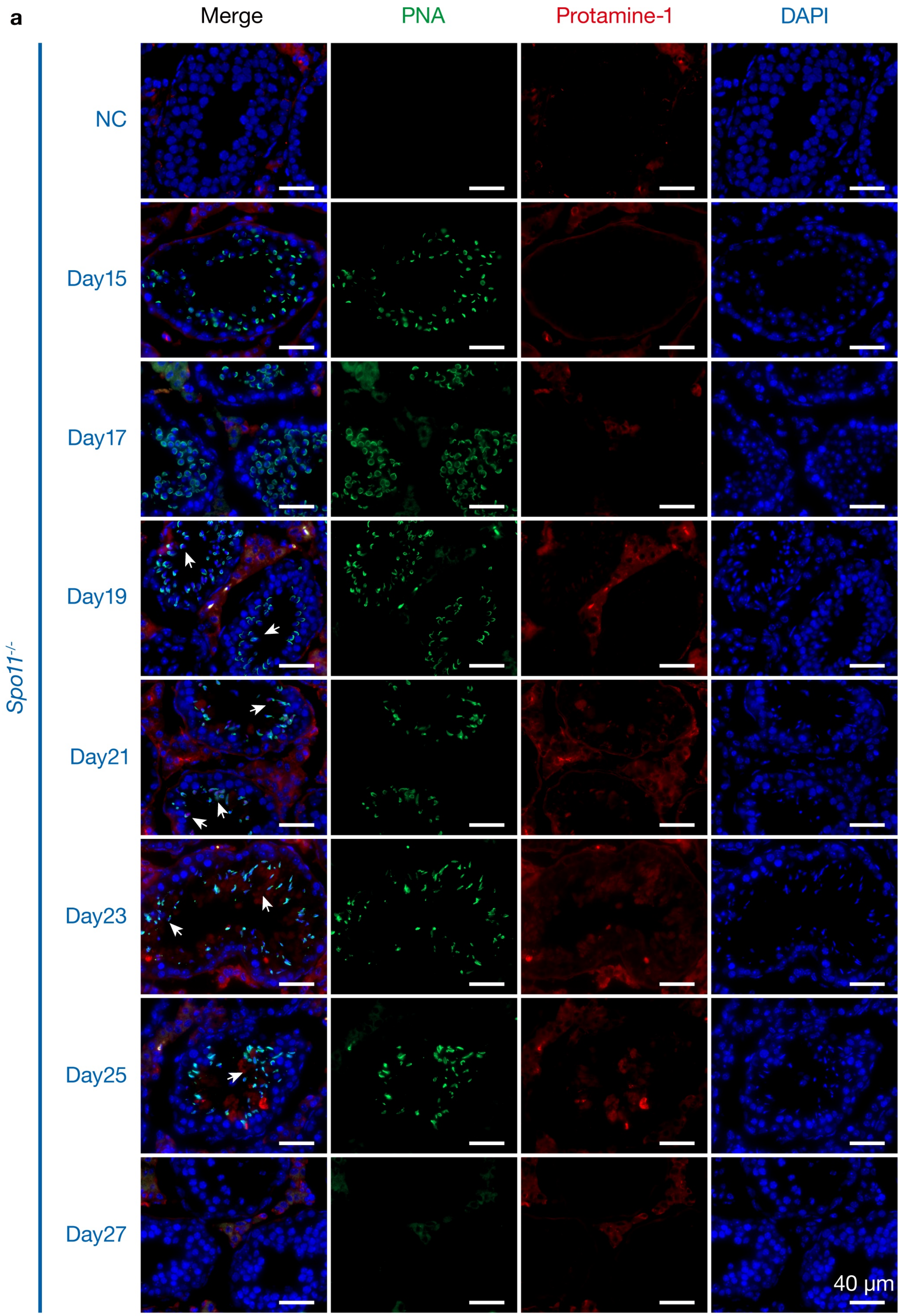

Jiang et al, Extended Data Fig. 6

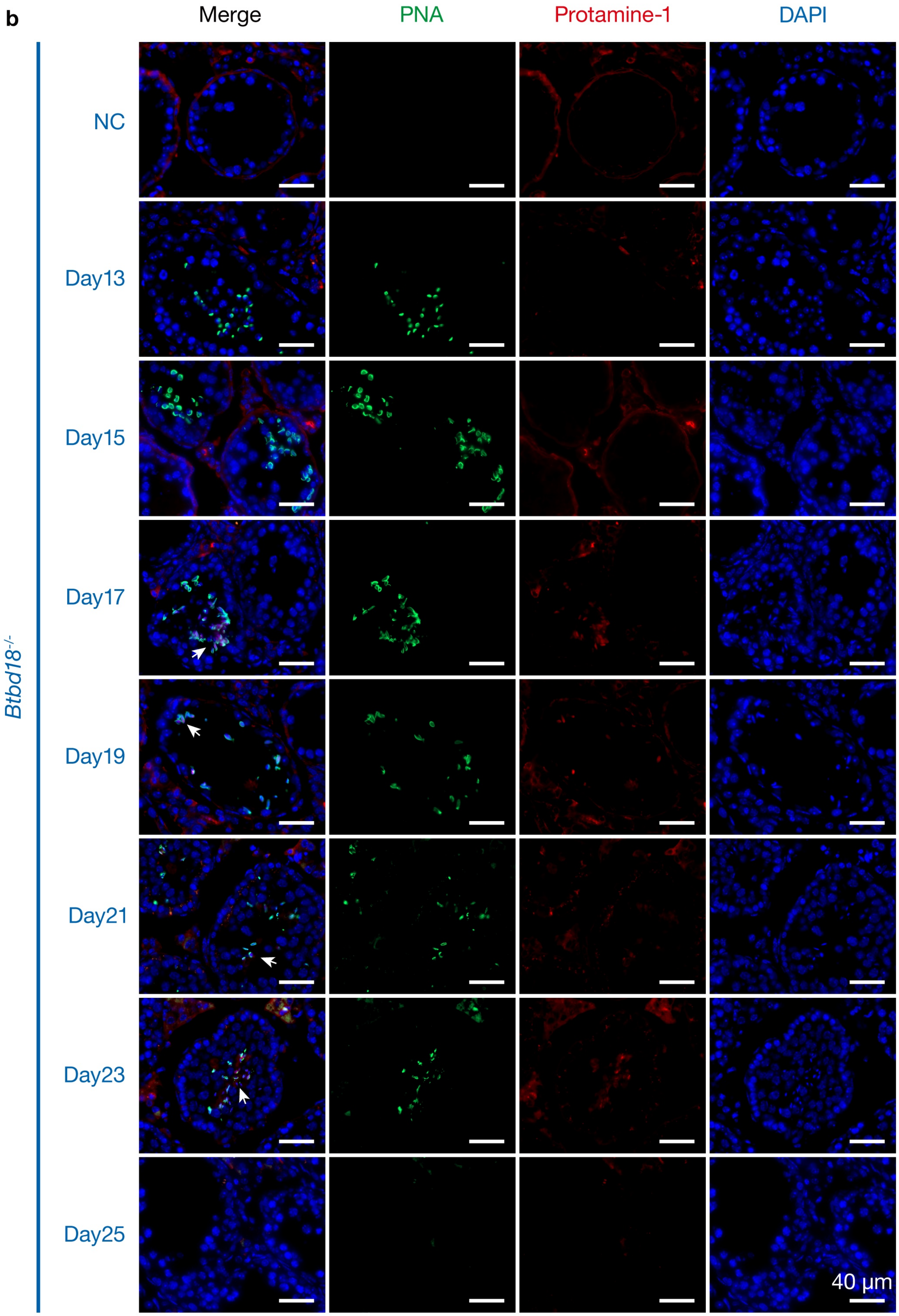

Jiang et al, Extended Data Fig. 6

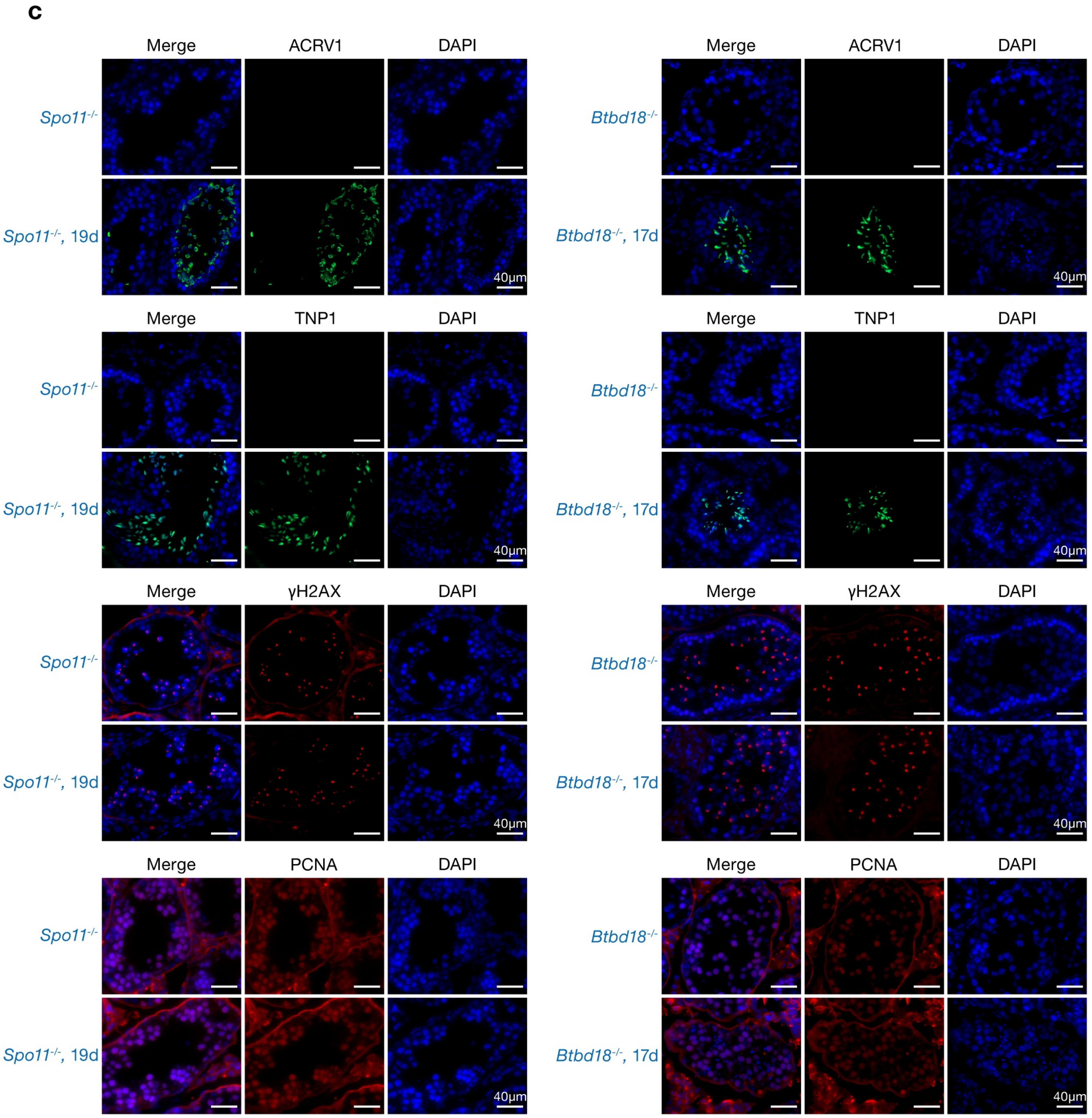

Jiang et al, Extended Data Fig. 6

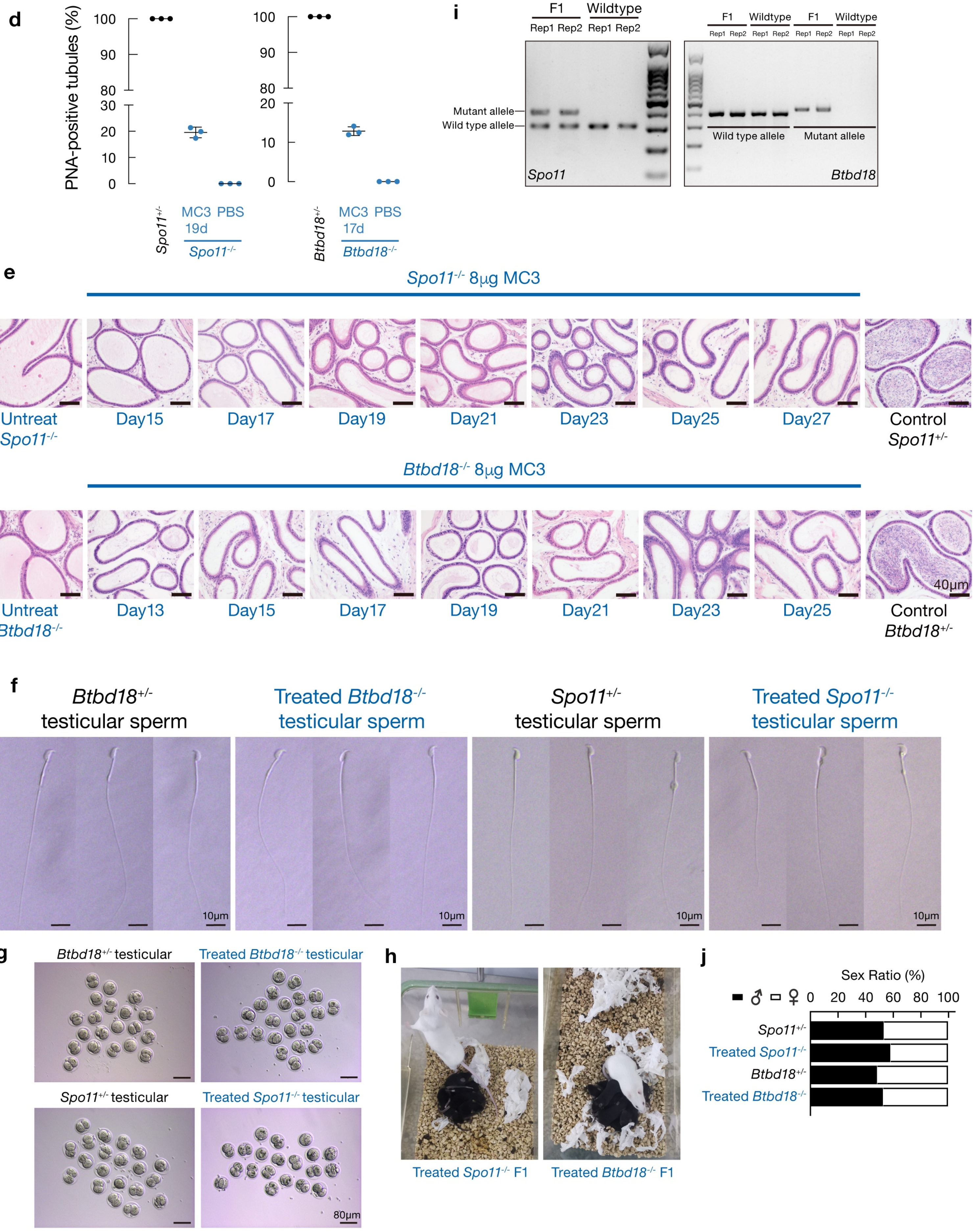

**Extended Data Figure 7. Systemic, epigenetic and genomic safety following mRNA–LNP treatment.**

**(a)** Histopathological assessment of major organs in *Papolb*$^{-/-}$, *Papolb*$^{-/-}$ treated (6 months), and *Papolb*$^{+/-}$ mice following intratubular injection of 8 µg MC3–*Papolb* mRNA. Scale bar, 50 µm. (n = 3 mice per group).

**(b–c)** Representative H&E images of major organs (heart, liver, spleen, lung, kidney) from treated *Papolb*$^{+/-}$ and *Papolb*$^{-/-}$ mice at day 1–21 post-treatment. Scale bar, 100 µm. (n = 3 mice per time point).

**(d)** Western blot of CYP17A1 with densitometric quantification normalized to GAPDH. Data are mean ± s.d. (n = 3 biological replicates). Statistical analysis: one-way ANOVA with Tukey's multiple comparisons test.

**(e)** Systemic safety evaluation by complete blood count and serum biochemistry. Serum biochemical analysis assessing liver enzymes (ALT, AST, ALP, T-Bil), kidney function (BUN, CRE), Glucose and LDH. Complete blood count Parameters shown include platelet count, RBC, HGB, HCT, MCV, MCH, MCHC, Neutrophil%, Lymphocyte%, Monocyte%, Basophil% and Eosinophil%. Data are mean ± s.d. (n = 5 mice per group per time point). Statistical analysis: two-way ANOVA with Tukey's multiple comparisons test. ALT, alanine aminotransferase; AST, aspartate aminotransferase; BUN, blood urea nitrogen; LDH, lactate dehydrogenase. ALP, alkaline phosphatase; T-Bil, total bilirubin; LDH, lactate dehydrogenase; RBC, red blood cell count; HGB, hemoglobin; PLT, platelet count; HCT, hematocrit; MCV, mean corpuscular volume; MCH, mean corpuscular hemoglobin; MCHC, mean corpuscular hemoglobin concentration.

**(f–g)** DNA methylation at imprinting DMRs (*Peg3*, *Snrpn*, *H19*) measured by pyrosequencing. Dots represent CpG sites; horizontal lines indicate mean methylation. Liver DNA was analyzed. Data are mean ± s.d. (Het = 3; Papi = 4). P-values for individual CpGs were adjusted using Benjamini–Hochberg correction; no CpG remained significant after correction. DMRs, differentially methylated regions; CpG, cytosinephosphate-guanine. *Peg3*, paternally expressed 3; *Snrpn*, small nuclear ribonucleoprotein polypeptide N; *H19*, H19 imprinted maternally expressed transcript.

**(h)** RNA-seq analysis of early embryos (n = 2 biological replicates per group) and adult tissues (n = 3 biological replicates per group) from Papi and *Papolb*$^{+/-}$ controls. Late-2-cell and 4-cell embryos and adult organs (heart, liver, lung, spleen, testis and kidney) were profiled. Differential expression analysis was performed using DESeq2; thresholds: FDR < 0.05 (genes highlighted in blue). Tpm, transcripts per million.

**(i)** Copy-number variation analysis using 1 Mb genomic bins comparing F0 and F1 individuals to control genomes. No large-scale deletions or insertions (>1 Mb) were detected (n = 3).

**(j)** PCR sensitivity analysis using serial dilutions of *Papolb* cDNA to establish detection limits for primer sets targeting *Papolb* mRNA; genomic DNA from offspring showed no amplification consistent with integration. (n = 3).

Jiang et al, Extended Data Fig. 7

**a**

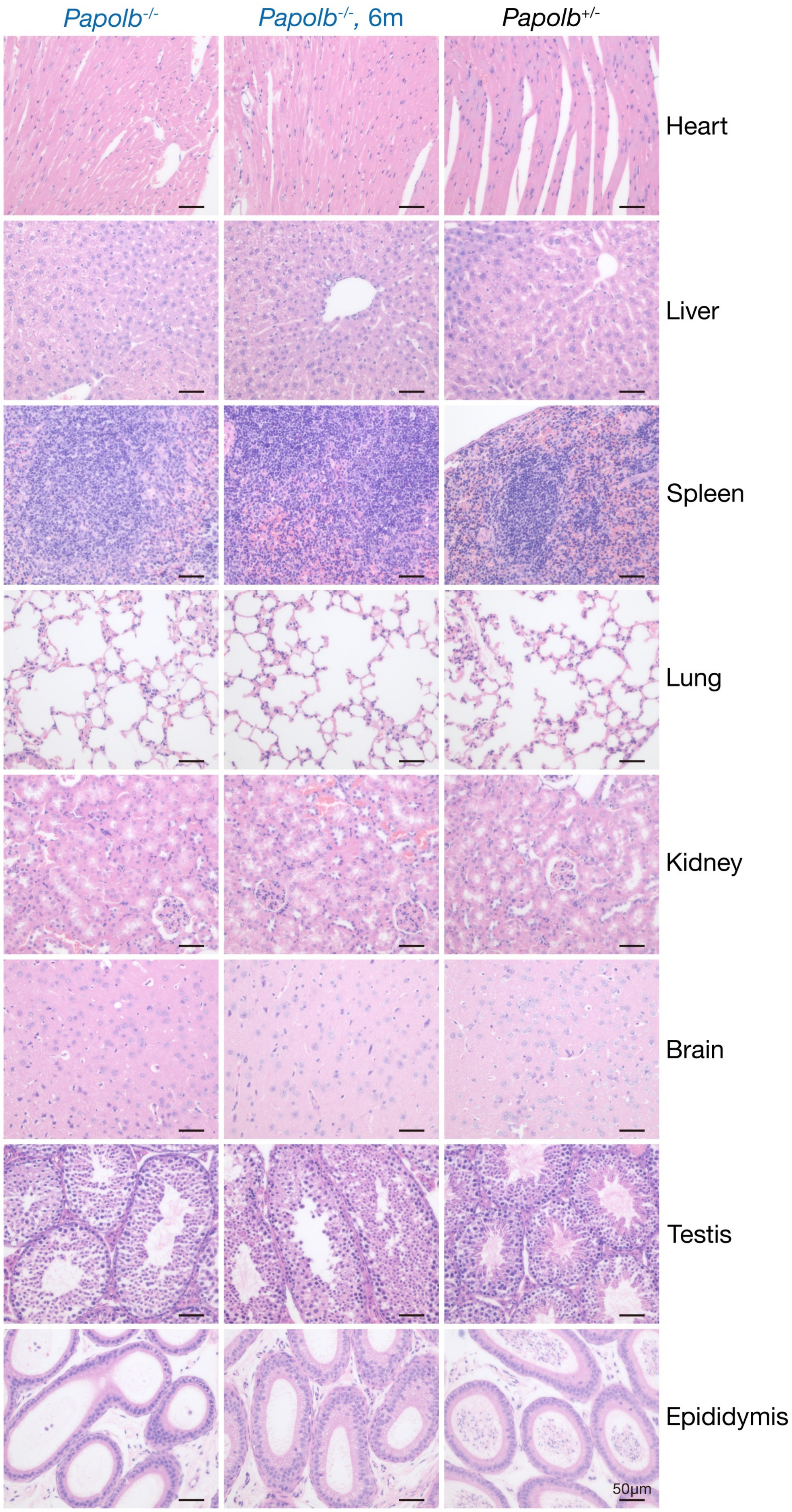

Jiang et al, Extended Data Fig. 7

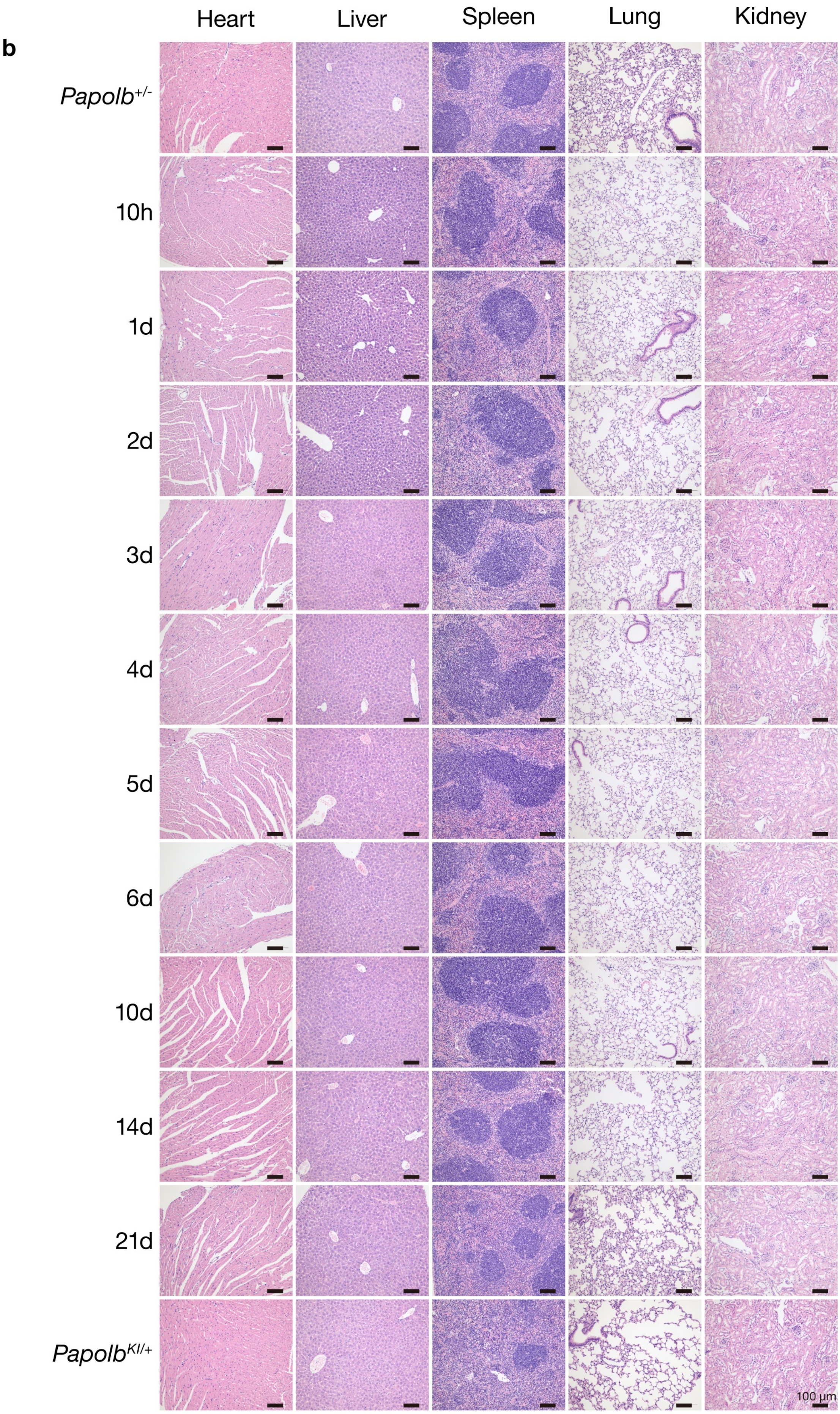

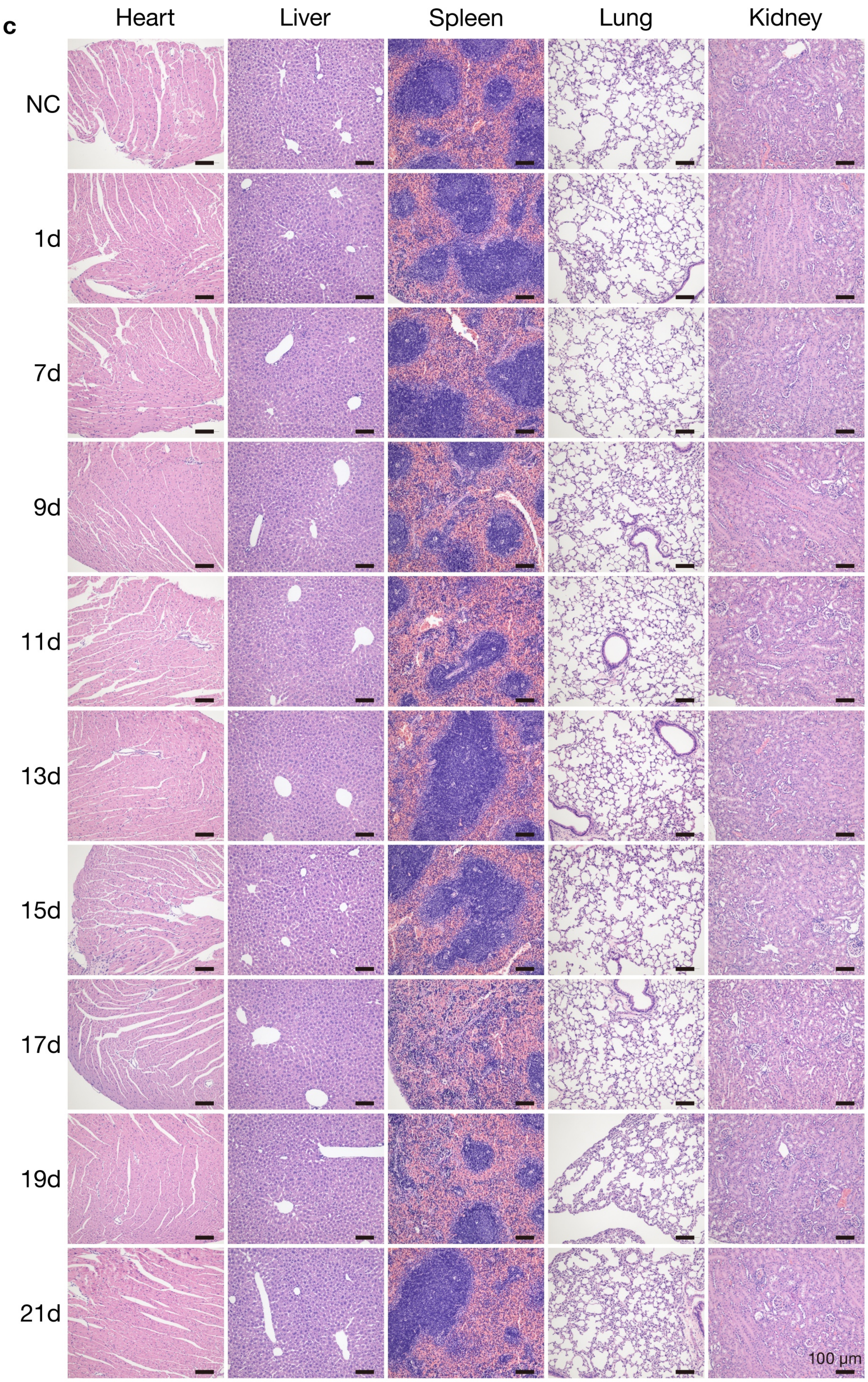
c
Heart
Liver
Spleen
Lung
Kidney
NC
1d
7d
9d
11d
13d
15d
17d
19d
21d
100 µm

Jiang et al, Extended Data Fig. 7

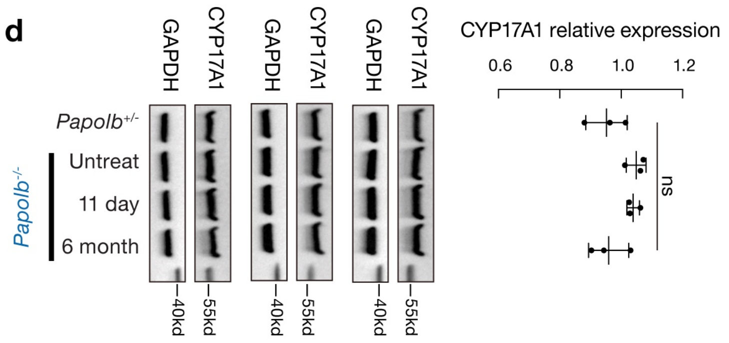


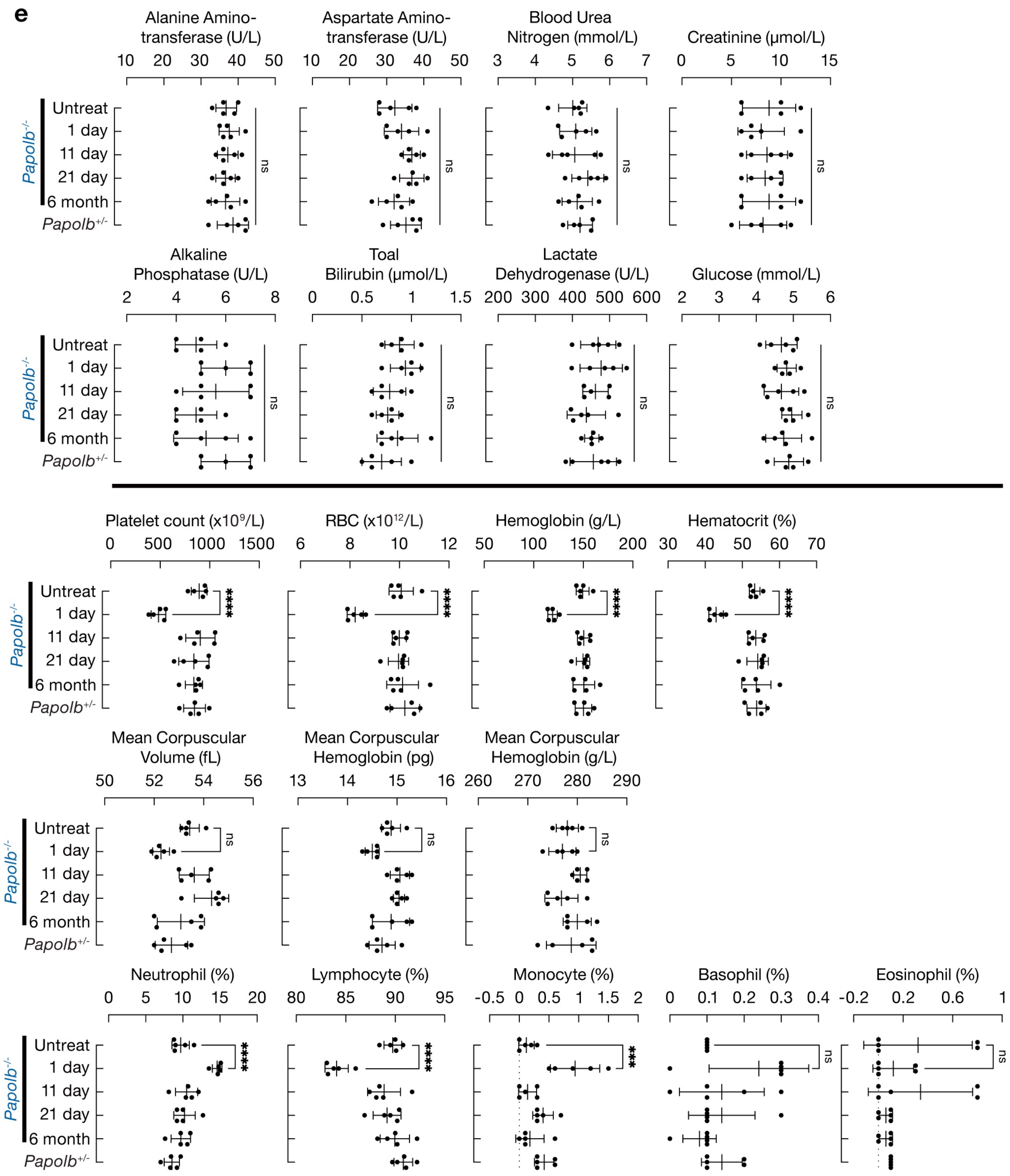

Jiang et al, Extended Data Fig. 7

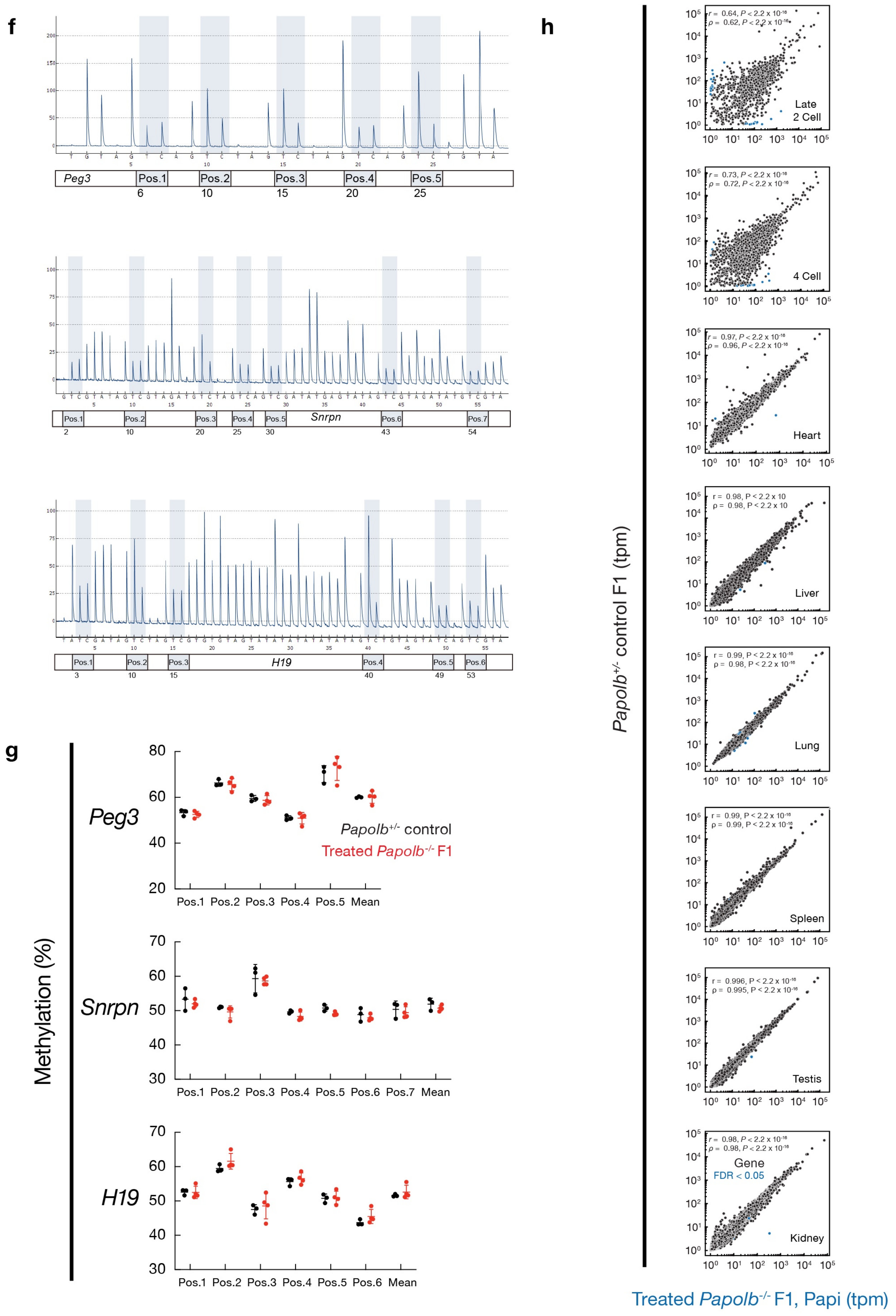

Jiang et al, Extended Data Fig. 7

i

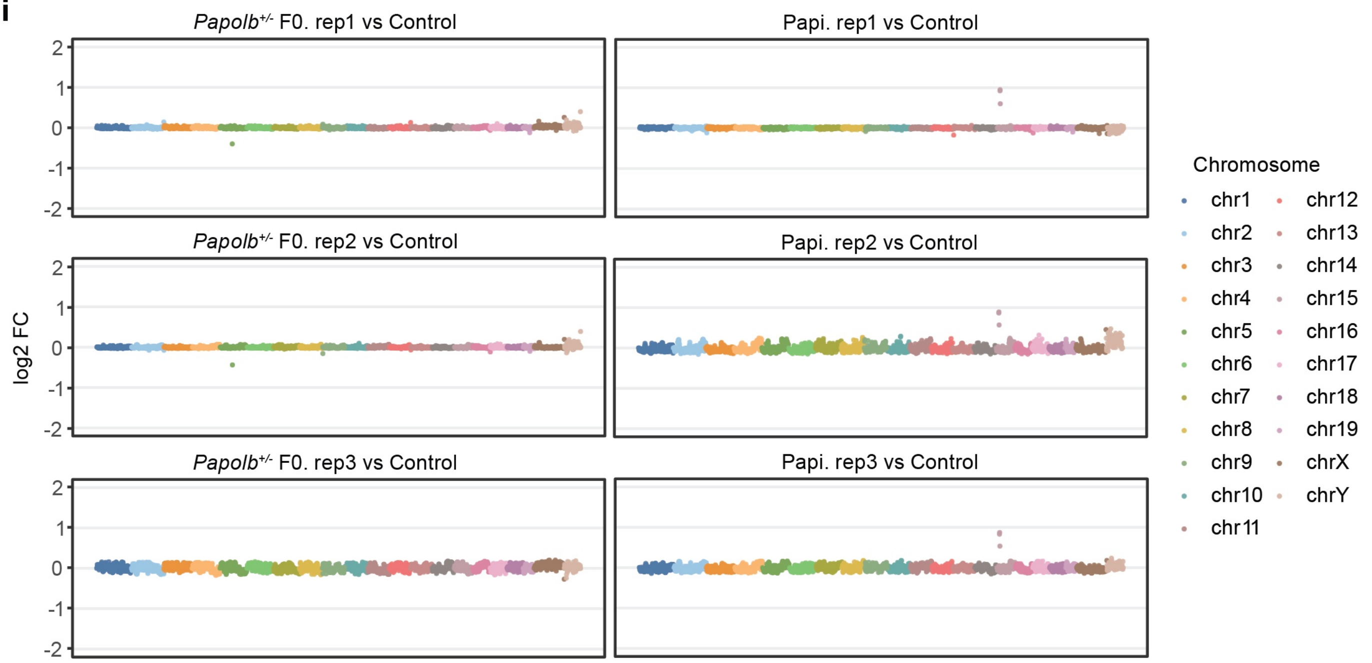


j

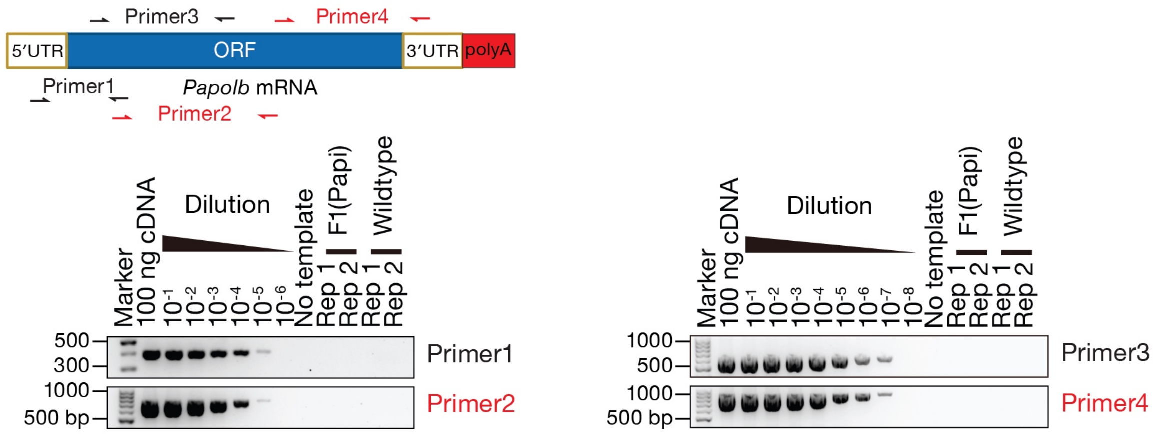

**Extended Data Figure 8. Efficient mRNA delivery and expression in human seminiferous tubules *ex vivo*.**

**(a)** Bright-field images of human (upper) and mouse (lower) seminiferous tubules 24 h after MC3–LNP incubation. Scale bars, 500 µm and 100 µm.

**(b)** Trypan blue staining assessing viability of human (left) and mouse (right) seminiferous tubules. Representative images from 3 donors and 6 mice. Scale bar, 100 µm.

**(c)** Quantification of luminescence from mouse seminiferous tubule cultures following LNP exposure. (n = 6 samples).

**(d)** Immunofluorescence staining for RNF17 (red) and HA (green) following *Papolb*-HA mRNA delivery in mouse tubule cultures. Scale bar, 20 µm. (n = 6 samples).

Jiang et al, Extended Data Fig. 8

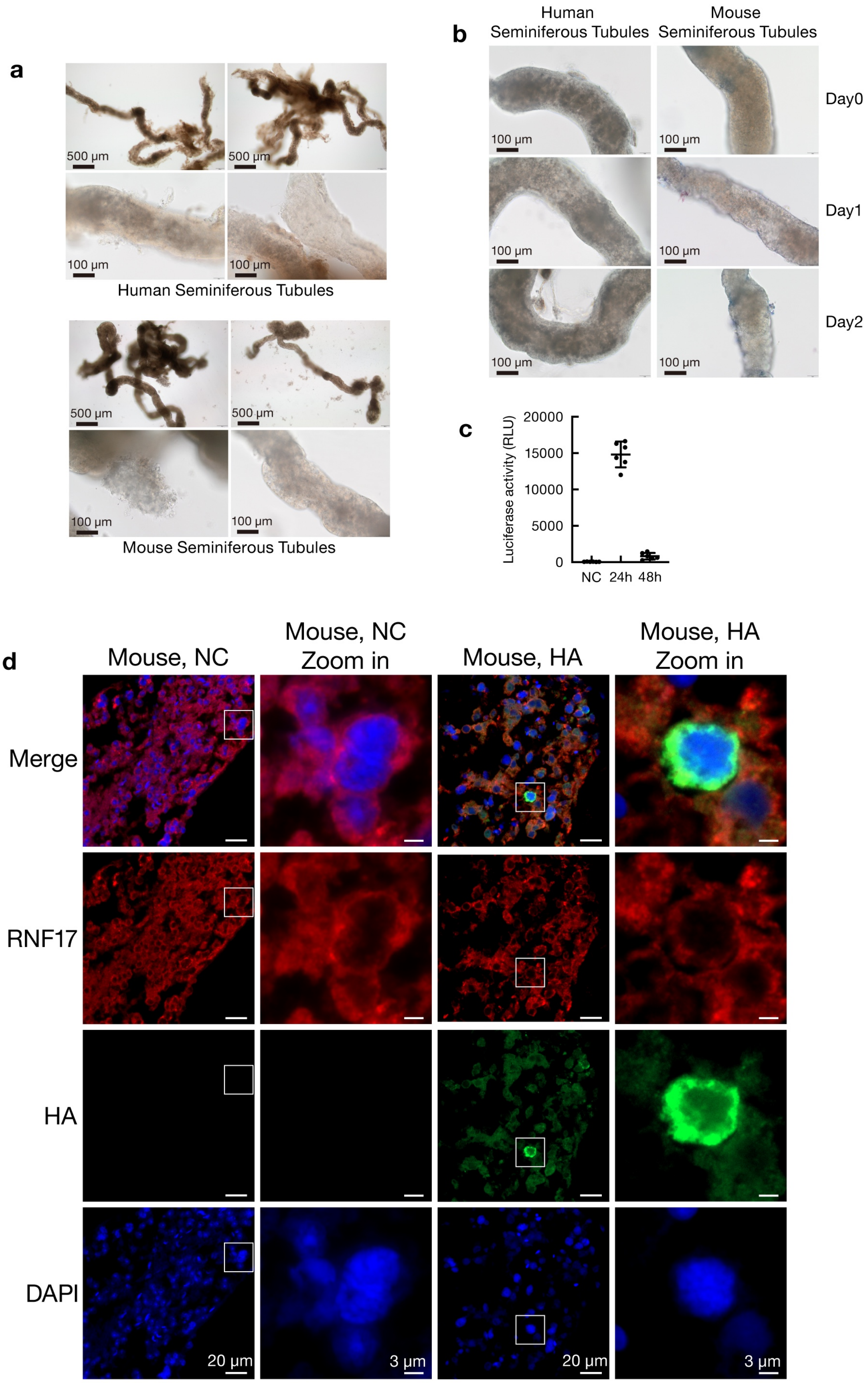
