## Extended Data Table1 for "Engineering temporal alignment of mRNA delivery enables functional rescue across genetic infertility models"

Supplementary Table 1: Sexual behavioral parameters of control and F1 (Papi) mice.

| | Control | F1 (Papi) | *p value* |
|---|---|---|---|
| N | 5 | 5 | |
| Mounting latency a | 63 ± 51 | 75 ± 78 | 0.442 |
| Intromission latency | 366 ± 278 | 446 ± 290 | 0.938 |
| Ejaculation latency | 134 ± 254 | 90 ± 99 | 0.102 |
| Post-ejaculatory mount latency | 279 ± 271 | 352 ± 137 | 0.215 |
| Duration of copulatory series | 40 ± 35 | 90 ± 41 | 0.765 |
| Number of mounts b | 12 ± 4 | 15 ± 5 | 0.794 |
| Number of intromissions | 15 ± 12 | 37 ± 20 | 0.310 |
| Number of copulatory series | 2 ± 1 | 3 ± 2 | 0.452 |

Mean ± s.d.; a unit is second (s). b unit is counts.
